# Immunoglobulin M regulates airway hyperresponsiveness independent of T helper 2 allergic inflammation

**DOI:** 10.1101/2023.08.02.551636

**Authors:** Sabelo Hadebe, Anca Flavia Savulescu, Jermaine Khumalo, Katelyn Jones, Sandisiwe Mangali, Nontobeko Mthembu, Fungai Musaigwa, Welcome Maepa, Hlumani Ndlovu, Amkele Ngomti, Martyna Scibiorek, Javan Okendo, Frank Brombacher

**Author notes:** **Corresponding Author**: Professor Frank Brombacher, International Centre for Genetic Engineering and Biotechnology (ICGEB) and Institute of Infectious Diseases and Molecular Medicine (IDM), Division of Immunology, Health Science Faculty, University of Cape Town, Cape Town, South Africa, Dr Sabelo Hadebe, Division of Immunology, Health Science Faculty, University of Cape Town, Cape Town, South Africa.

## Abstract

Allergic asthma is a disease driven by T helper 2 (Th2) cells, eosinophilia, airway hyperresponsiveness (AHR) and IgE-secreting B cells. Asthma is largely controlled by corticosteroids and β_2_ adregenic receptor agonists that target and relax airway smooth muscle (ASM). Immunoglobulin M (IgM) isotype secreted by naïve B cells is important for class switching but may have other undefined functions. We investigated the role of IgM in a house dust mite (HDM)-induced Th2 allergic asthma model. We sensitised wild-type (WT) and IgM-deficient (IgM KO) mice with HDM and measured AHR, and Th2 responses. We performed RNA sequencing on the whole lung of WT and IgM KO mice sensitised to saline or HDM. We validated our AHR data on human ASM by deleting genes using CRISPR and measuring contraction by single-cell force cytometry. We found IgM to be essential in AHR but not Th2 airway inflammation or eosinophilia. RNA sequencing of lung tissue suggested that IgM regulated AHR through modulating brain-specific angiogenesis inhibitor 1-associated protein 2-like protein 1 (*Baiap2l1*) and other genes. Deletion of *BAIAP2L1* led to a differential reduction in human ASM contraction when stimulated with TNF-α and Acetylcholine, but not IL-13. These findings have implications for future treatment of asthma beyond current therapies.

## Introduction

Immunoglobulin M (IgM) is the first antibody isotype expressed during B cell development and the first humoral antibody responder, conserved across all species from zebrafish to humans (Akula et al., 2014). IgM can be divided into natural and antigen-induced IgM and can either be membrane-bound IgM-type B cell receptor (BCR) or secreted IgM (Baumgarth et al., 2000; Blandino and Baumgarth, 2019). Natural IgM plays multiple roles in homeostasis including scavenging and clearance of apoptotic cell debris in conjunction with phagocytic macrophages, B cell survival through tonic signals, lymphoid tissue architecture and prevention of autoimmune diseases (Ehrenstein and Notley, 2010; Michaud et al., 2020; Quartier et al., 2005). At mucosal sites, both natural and antigen-induced IgM play a role in shaping healthy microbiota and IgM repertoire is also shaped by microbiota (New et al., 2020; Wesemann et al., 2013). Secreted IgM antigen complexes can connect signals via unique and shared receptors, suggestive of a more pleiotropic role in homeostasis and disease states (Jones et al., 2020; Kawahara et al., 2003; Nguyen et al., 2017a).

Natural IgM and secreted IgM are essential in many infectious and non-infectious diseases including those induced by parasites, fungi, bacterial, viral and autoimmune diseases (Jones et al., 2020). In *Plasmodium falciparum*, anti-α-gal IgM and anti-MSP1 directed antibodies are protective against primary and secondary infections (Krishnamurty et al., 2016; Yilmaz et al., 2014). Mice deficient of secreted IgM are susceptible to pulmonary *Cryptococcus neoformans* and *P. carinii* infection partly due to reduced activation of innate and adaptive responses (Rapaka et al., 2010; Subramaniam et al., 2010). At mucosal surfaces, IgM promotes healthy gut bacteria that is beneficial for homeostasis such as Firmicutes and Bacteroidetes (Magri et al., 2017). Natural and induced secreted IgM produced mainly by B1a cells is protective against *Streptococcus pneumoniae* and *Francisella tularensis* infection (del Barrio et al., 2015; Weber et al., 2014). In these settings, the protective effects of sIgM depended on cytokines IL-1β and GM-CSF (del Barrio et al., 2015; Weber et al., 2014). The involvement of natural or induced IgM in allergic asthma is unknown, despite selective IgM syndrome dominated by asthma patients (Goldstein et al., 2006).

Asthma is a T helper 2 (Th2) disease characterised by eosinophilic lung inflammation, mucus production, airway hyperresponsiveness (AHR), Th2 cytokines (interleukin-4 (IL-4), IL-5 and IL-13) and B cells producing IgE (Lambrecht and Hammad, 2014). IgM is central to class switch recombination that results in IgE class-switched B cells (Mandler et al., 1993). We and others recently showed that the role of B cells in asthma is complex, where the load of the antigen is crucial in their function (Dullaers et al., 2017; Habener et al., 2021; Hadebe et al., 2021). Mice deficient of B cells (μMT KO) can mount exaggerated AHR when challenged with house dust mite (HDM) (Ballesteros-Tato et al., 2016; Dullaers et al., 2017; Wypych et al., 2020) or ovalbumin (OVA) (Hamelmann et al., 1999; Korsgren et al., 1997; MacLean et al., 1999) partly due to lack of regulatory B cells that dampen AHR (Habener et al., 2021). Furthermore, interleukin 4 receptor alpha signalling in B cells is required for optimal Th2 allergic airway inflammation through regulation of AHR, germinal centre (GC) formation and B effector 2 function (Hadebe et al., 2021).

Interestingly, B cell isotypes show unique functions in allergic asthma, for example, IgE and its high-affinity receptor, FcεR are redundant in an HDM model (McKnight et al., 2017), whereas IgD plays an amplifying and regulatory role in various allergic models (Shan et al., 2018). In this context, IgD activates basophils to secrete IL-4 and Th2 induction during the sensitisation stage through binding basophils via galectin-9 and CD44. Once Th2 responses have been amplified, IgD ligation blocked IgE-mediated basophil degranulation through competing for antigen and inhibiting FcεR mediated signalling (Shan et al., 2018). How IgM isotype contributes in the development of allergic asthma is unclear. It is also unclear whether secreted IgM plays different roles compared to membrane-bound IgM, which is more likely to undergo class switching to IgE. We challenged mice lacking both membrane and secreted IgM with HDM and other allergens and found a profound reduction in AHR. RNA sequencing of lung tissue showed a downregulation of brain-specific angiogenesis inhibitor 1-associated protein 2-like protein 1 (*Baiap2l1*) and erythroid differentiation regulatory factor 1 (*Erdr1*) genes associated with actin cytoskeleton and rearrangement smooth muscle contraction. As a proof of principle, we showed in human smooth muscle cell line that deletion of these genes via CRISPR-resulted in a reduction in smooth muscle contraction at a single cell level. These are unexpected functions of secreted, and membrane-bound IgM; namely, its involvement in modulating airway smooth muscle contraction.

## Results

### IgM-deficient mice show profound airway hyperresponsiveness reduction when exposed to HDM

We sensitised and challenged IgM-deficient (IgM KO) and wild-type Balb/c (WT) mice with HDM intratracheally (i.t.) and analysed AHR, lung infiltrates and cytokines (Figure 1A, Figure 1-figure supplement 1A). We found moderately reduced resistance and elastance in IgM KO sensitised with a high dose of HDM (100μg) compared to WT mice (Figure 1-figure supplement 1B). Similarly, we also observed a profound reduction in AHR in IgM KO mice sensitised with a low dose of HDM (1μg) compared to WT (Figure 1B). Interestingly, lung eosinophils were intact at both low dose (Figure 1C) and high dose HDM (Figure 1-figure supplement 1D). We could also show that AHR reduction in IgM KO mice was not allergen-specific, as we observed similar findings using ovalbumin (OVA) complexed to alum adjuvant (Figure 1-figure supplement 2A-B) and acute papain-induced allergic inflammation (Figure 1-figure supplement 2C-D). We observed similar finding of profound AHR reduction in IgM KO mice in C57BL/6 background compared to WT mice (Figure 1 - figure supplement 3A). These findings suggested that IgM-deficient mice have profound AHR reduction independent of allergen or mice background strain.

**Figure 1.**
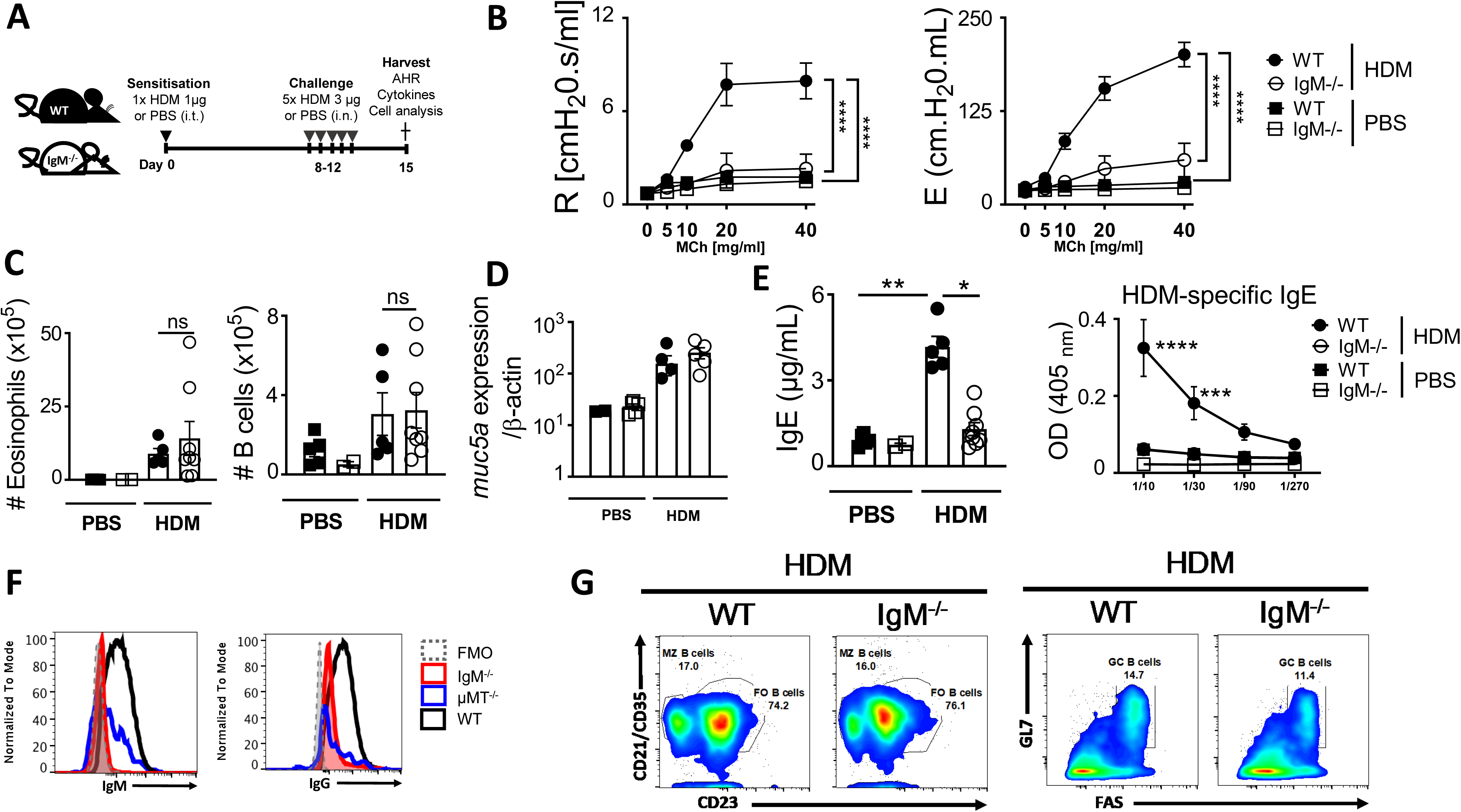
lgM-deficiency leads to reduced airway hyperresponsiveness and class switching to IgE in HDM-induced asthma. **(A)** Schematic diagram showing sensitisation and challenge protocol where mice (IgM KO) and wild type littermate control (WT) were sensitised with HDM 1 μg intra-tracheally on days 0 and challenged with HDM 3 μg on days 8-12. Analysis was done on day 15. **(B)** Airway resistance and elastance were measured with increasing doses of acetyl methacholine (0 −40 mg/mL). **(C)** Total lung eosinophil numbers (live^+^Siglec-F^+^CD11c^-^) and B cells (live^+^B220^+^CD19^+^MHCII^+^) were stained and analysed by Flow cytometry and enumerated from % of live cells. **(D)** *Muc5a* gene expression in whole lung tissue. **(E)** Total IgE and HDM-specific IgE in serum. **(F)** IgG1 and IgM surface expression in mediastinal lymph node B cells of WT, IgM KO and μMT KO mice. **(G)** Marginal Zone (live^+^B220^+^CD19^+^MHCII^+^CD21/CD35^+^CD23^-^), follicular (live^+^B220^+^CD19^+^MHCII^+^CD23^+^CD21/CD35^+^) and Germinal Centre (live^+^B220^+^CD19^+^MHCII^+^GL7^+^FAS^+^) B cells in the mediastinal lymph node of WT and IgM KO mice challenged with HDM. Shown is mean ± SEM from two pooled experiments (n=7 - 10). Significant differences between groups were performed by Student *t*-test (Mann-Whitney) (c, d, e) or by Two-Way ANOVA with Benforroni post-test (b) and are described as: \**p*<0.05, \*\**p*<0.01, \*\*\**p*<0.001, \*\*\*\**p*<0.0001.

### IgM deficiency does not impact B cell subsets in primary and secondary lymphoid organs, but class switching is impaired

We also observed no significant changes between IgM KO and WT mice in mucus production shown by *Muc5a* gene expression (Figure 1D) or goblet cells in Balb/C (Figure 1 - figure supplement 1F) and C57BL/6 mice challenged with HDM (Figure 1-figure supplement 3B). Accompanying reduction in AHR in IgM KO mice was low titers of total IgE, HDM-specific IgE, and mediastinal lymph node (mLN) B cell surface expression of IgM and IgG1 (Figure 1E-F), owing to a lack of class switching. Interestingly, the numbers of lung B cells were normal (Figure 1C and Figure 1 - figure supplement 1C) and the frequencies of B cell subsets such as follicular, marginal zones and germinal centres (GCs) B cells were not affected by lack of IgM (Figure 1G and Figure 1-figure supplement 1G). The lack in class switching in Balb/C mice was also consistent with what we found in IgM KO mice in the C57BL/6 background (Figure 1-figure supplement 3C). This suggested a normal interaction between B and T cells in GCs, but lack of AID-dependent class switching, despite increased expression of IgD (Figure 1-figure supplement 4A) (Muramatsu et al., 2000). We checked for natural IgM and antigen-induced IgM in multiple tissues. B cells expressing IgD were increased in all tissues including mLNs, peritoneal cavity and spleen (Figure 1-figure supplements 4A-C) and pre-B cell subsets were normal in BM (Figure 1-figure supplement 4D) in the absence of IgM as previously reported (Lutz et al., 1998).

### T helper 2 cells are intact in the absence of IgM, and serum transfer can partially restore antibody function

To investigate whether the reduction in AHR was due to reduced Th2 cells and cytokines, we stimulated total mLN and lung cells with anti-CD3 for 5 days or with PMA/ionomycin for 5 hrs in the presence of monensin and measured secreted or intracellular IL-4 and IL-13 expression. We found no differences in CD4 T cells and T follicular helper cells (Figure 2A) in mLN and in secreted or intracellular levels of IL-4 and IL-13 between IgM KO and WT mice challenged with HDM in both mLN and lungs (Figure 2B-C). To investigate what drives this reduced AHR in the absence of IgM between secreted antibodies or the IgM B cell receptor on the surface of B cells, we transferred serum from naïve WT mice into IgM KO mice a day before sensitization, during sensitisation and a day before challenge with HDM allergen (Figure 2D), as previously described (Wojciechowski et al., 2009). AHR in IgM KO was still reduced compared to WT mice even after the transfer of WT serum (Figure 2E), but levels of total IgE, HDM-specific IgE and HDM-specific IgG1 were increased and comparable to those found in WT mice (Figure 2F and Figure 2-figure supplement 1D). The redundant role of IgE was consistent with previous studies where IgE nor its high-affinity receptor (FcεRI) were essential in AHR in an HDM or OVA models (McKnight et al., 2017). The lack of functional role of serum-transferred IgE was consistent with earlier findings on *H. polygyrus* transfer of immune serum where IgE was found not to be essential in protection against *H. polygyrus* re-infection (Wojciechowski et al., 2009).

**Figure 2.**
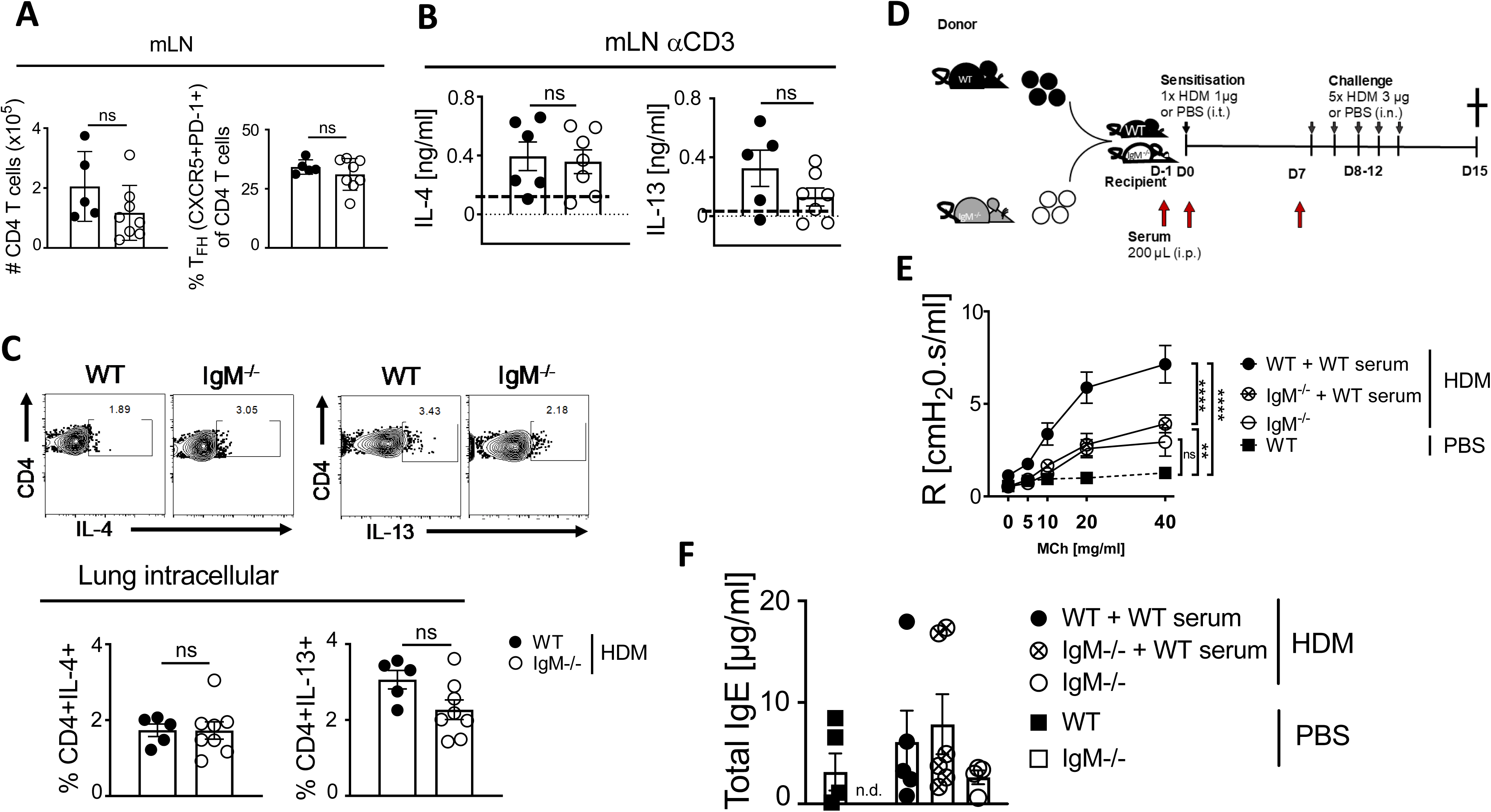
lgM-deficiency does not lead to reduced Th2 allergic airway inflammation and serum transfer restores IgE, but not AHR. Fig. a-c, mice treated as in Figure 1A. **(A)** Total mediastinal lymph node CD4 T cell numbers (live^+^CD3^+^CD4^+^) and % of Follicular Helper T cells (live^+^CD3^+^CD4^+^PD-1^+^CXCR5^+^) of CD4 T cells were stained and analysed by Flow cytometry and enumerated from % of live cells. **(B)** Mediastinal lymph nodes were stimulated with anti-CD3 (10μg/mL) for 5 days and supernatants were used to measure levels of IL-4 and IL-13. Cytokines were not detected in unstimulated or HDM (30μg) stimulated mLN. **(C)** Representative FACS plots and frequencies of lung CD4 T cells (live^+^CD3^+^CD4^+^) producing IL-4 and IL-13 after 5 hr stimulation with PMA/ionomycin in the presence of monensin. **(D)** Schematic diagram showing serum transfer from WT to IgM KO which were then sensitised as shown in Figure 1,A. **(E)** Airway resistance was measured with increasing doses of acetyl methacholine (0-40 mg/mL). **(F)** Total IgE in serum of mice either transferred with WT serum, IgM KO serum or no serum. Shown is the mean ± SEM from two pooled experiments (n=5 - 8). Significant differences between groups were performed by Student *t*-test (Mann-Whitney) (C, D, E) or by Tw*o-Way ANOVA with Benforroni post-test (B) and are described as: *p<0.05, \*\**p<0.01, \*\*\**p*<0.001, \*\*\*\**p*< 0.0001.

### Replacement of IgM-deficient mice with functional hematopoietic cells in busulfan mice chimeric mice restores airway hyperresponsiveness

We then generated bone marrow chimeras by chemical radiation using busulfan (Montecino-Rodriguez and Dorshkind, 2020). We treated mice three times with busulfan for 3 consecutive days and after 24 hrs transferred naïve bone marrow from congenic CD45.1 WT mice or CD45.2 IgM KO mice (Figure 3A and Figure 3-figure supplement 1A). We showed that recipient mice that did not receive donor bone marrow after 4 days post-treatment had significantly reduced lineage markers (CD45^+^Sca-1^+^) or lineage negative (Lin^-^) cells in the bone marrow when compared to untreated or vehicle (10% DMSO) treated mice (Figure 3-figure supplements 1B-C). We allowed mice to reconstitute bone marrow for 8 weeks before sensitisation and challenge with low dose HDM (Figure 3A). We showed that WT (CD45.2) recipient mice that received WT (CD45.1) donor bone marrow had higher airway resistance and elastance and this was comparable to IgM KO (CD45.2) recipient mice that received donor WT (CD45.1) bone marrow (Figure 3B). As expected, IgM KO (CD45.2) recipient mice that received donor IgM KO (CD45.2) bone marrow had significantly lower AHR compared to WT (CD45.2) or IgM KO (CD45.2) recipient mice that received WT (CD45.1) bone marrow (Figure 3B). We confirmed that the differences observed were not due to differences in bone marrow reconstitution as we saw similar frequencies of CD45.1 cells within the lymphocyte populations in the lungs and other tissues (Figure 3-figure supplement 1D). We observed no significant changes in the lung neutrophils, eosinophils, inflammatory macrophages, CD4 T cells or B cells in WT or IgM KO (CD45.2) recipient mice that received donor WT (CD45.1/CD45.2) or IgM KO (CD45.2) bone marrow when sensitised and challenged with low dose HDM (Figure 3C).

**Figure 3.**
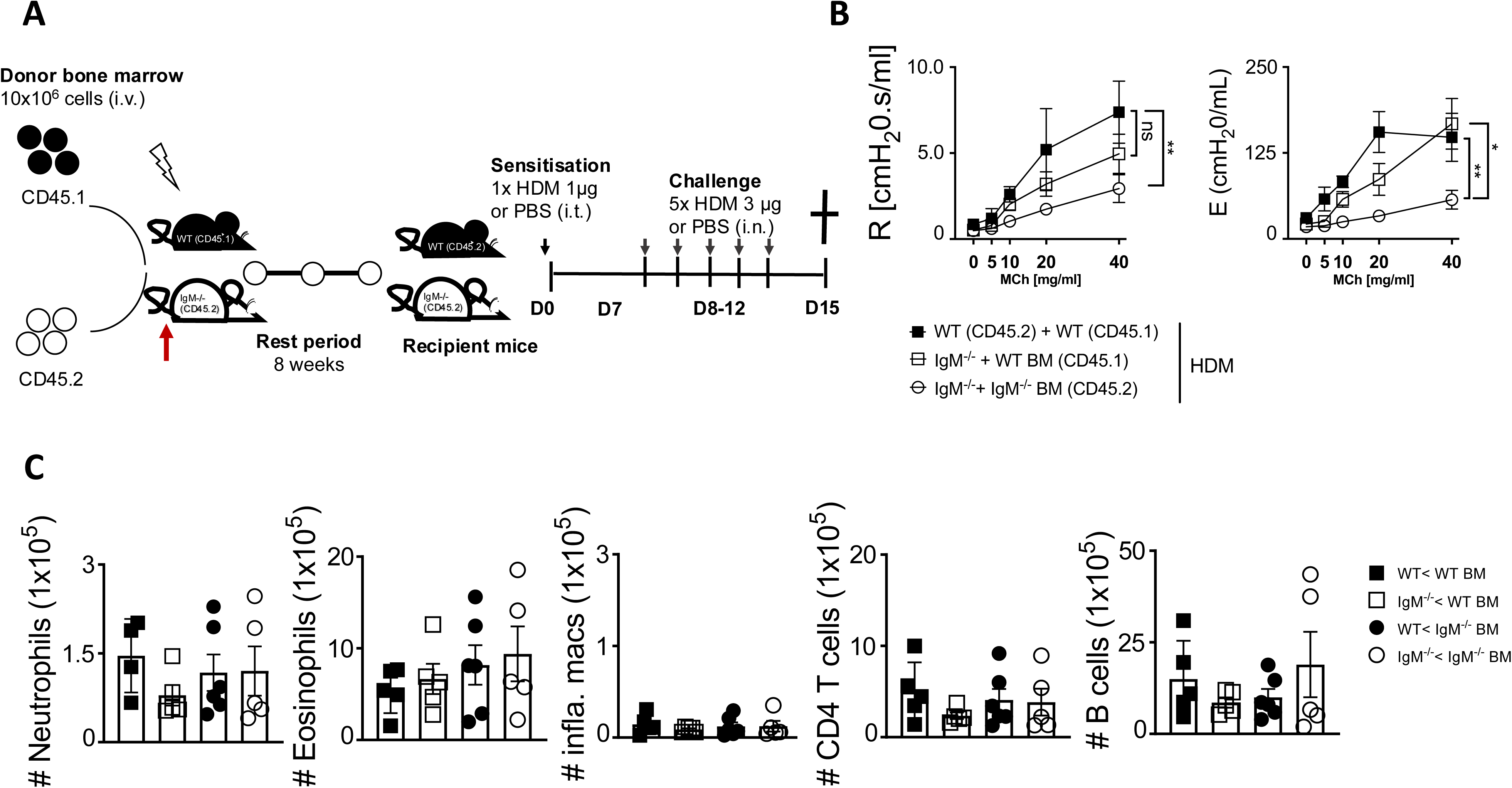
Partial wild type bone marrow replenishment restores AHR in IgM-deficient mice (Figure A-C, mice treated as in Figure 1A). **(A)** Schematic diagram showing WT and IgM-deficient mice being chemically irradiated with busulfan (25 mg per day for 3 days) and adoptively transferred with congenic bone marrow (10×10^6^ per mouse intravenously) at day 4. Mice (WT to IgM KO plus bone marrow) were then rested for 8 weeks before being sensitised as shown in Figure 1,A. **(B)** Airway resistance and elastance were measured with increasing doses of acetyl methacholine (0 −40 mg/mL). **(C)** Total lung neutrophils (live^+^CD11b^+^Ly6G^+^), eosinophils (live^+^Siglec-F^+^CD11c^-^), inflammatory macrophages (live^+^Ly6G^-^CD11b^+^F4/80^+^), CD4 T cells (live^+^CD3^+^CD4^+^CD8^-^) and B cells (live^+^B220^+^CD19^+^MHCII^+^) were stained and analysed by Flow cytometry and enumerated from % of live cells. Shown is the mean ± SD from one experiments (n=5 – 6 per group). Significant differences between groups were performed by Student *t*-test (Mann-Whitney) (C) or by Tw*o-Way ANOVA* with Benforroni post-test (B) and are described as: *p<0.05, *\*\**p<0.01, \*\*\**p*<0.001, \*\*\*\**p*< 0.0001.

Restoring IgM function through adoptive reconstitution with congenic CD45.1 bone marrow in non-chemically irradiated recipient mice or sorted B cells into IgM KO mice (Figure 2-figure supplement 1A) did not replenish IgM B cells to levels observed in WT mice and as a result did not restore AHR, total IgE and IgM in these mice (Figure 2-figure supplements 1B-C).

IgM is known to play key role in shaping gut microbiota, and its diversity can be shaped by gut commensals (Magri et al., 2017; Smith et al., 2023). To understand whether microbiota could influence AHR in IgM-deficient mice, we treated WT and IgM KO with an antibiotic cocktail 3x per week for 2 weeks (Figure 3-figure supplement 2A). We confirmed the reduction of bacteria in the faecal pellet after 2 weeks of antibiotic treatment (Figure 3-figure supplement 2A). We then sensitised and challenged these mice with HDM and measure AHR (Figure 3-figure supplement 2B). We observed increased resistance and elastance in WT mice compared to IgM KO mice and pre-treatment with antibiotics did not influence AHR (Figure 3-figure supplement 2B). Treatment with antibiotics did increase total IgE, HDM-specific IgE and IgG1 in WT mice as expected (Trompette et al., 2014), but had no impact on IgM-deficient mice (Figure 3-figure supplement 2C), suggesting that overall antibiotics did not influence AHR.

### RNA sequencing reveals enrichment of genes associated with skeletal muscle contraction and actin re-arrangement

Because we had found no other changes in asthmatic allergic features between WT and IgM KO, except for profoundly reduced AHR which we could only restore when we partially replaced IgM-deficient haematopoietic cells with WT bone marrow, we resorted to RNA sequencing. We wanted to know if there were lung-specific factors that influenced AHR regulation in IgM-deficient mice. Principal component analysis (PCA) depicted distinct global transcriptional changes with PC1 and PC2 explaining most of the variation (Figure 4A). We found a smaller variation in gene expression between WT and IgM KO in both HDM-challenged and PBS control mice (Figure 4B). We could mainly detect downregulation of genes such as brain-specific angiogenesis inhibitor 1-associated protein 2-like protein 1 (*Baiap2l1*), erythroid differentiation regulatory factor 1 (*Erdr1*), chemokines such as *Ccl8*, *Ccl9*, *Ccl17* and *Ccl22* in IgM KO mice challenged with HDM (Figure 4C-D). Interestingly, *Baiap2l1* and *Erdr1* were also downregulated in IgM KO saline-treated control mice, suggesting an inflammation-independent effect (Figure 4C and E). We also found the presence of the J-chain coding gene in WT mice which was absent in IgM KO mice, confirming a deletion in genes associated with holding IgM monomers together and thus IgM (Norderhaug et al., 1999) (Figure 4E). Gene set enrichment analyses (GSEA) confirmed that the genes contributing to changes in AHR between WT and IgM KO were associated with muscle system processes and skeletal muscle contraction (Figure 4F). There was also an over-representation of gene ratios associated with skeletal muscle development, differentiation and contraction, and suppression of genes associated with chemotaxis and plasma membrane-bounded cell projection (Figure 4-figure supplement 1).

**Figure 4.**
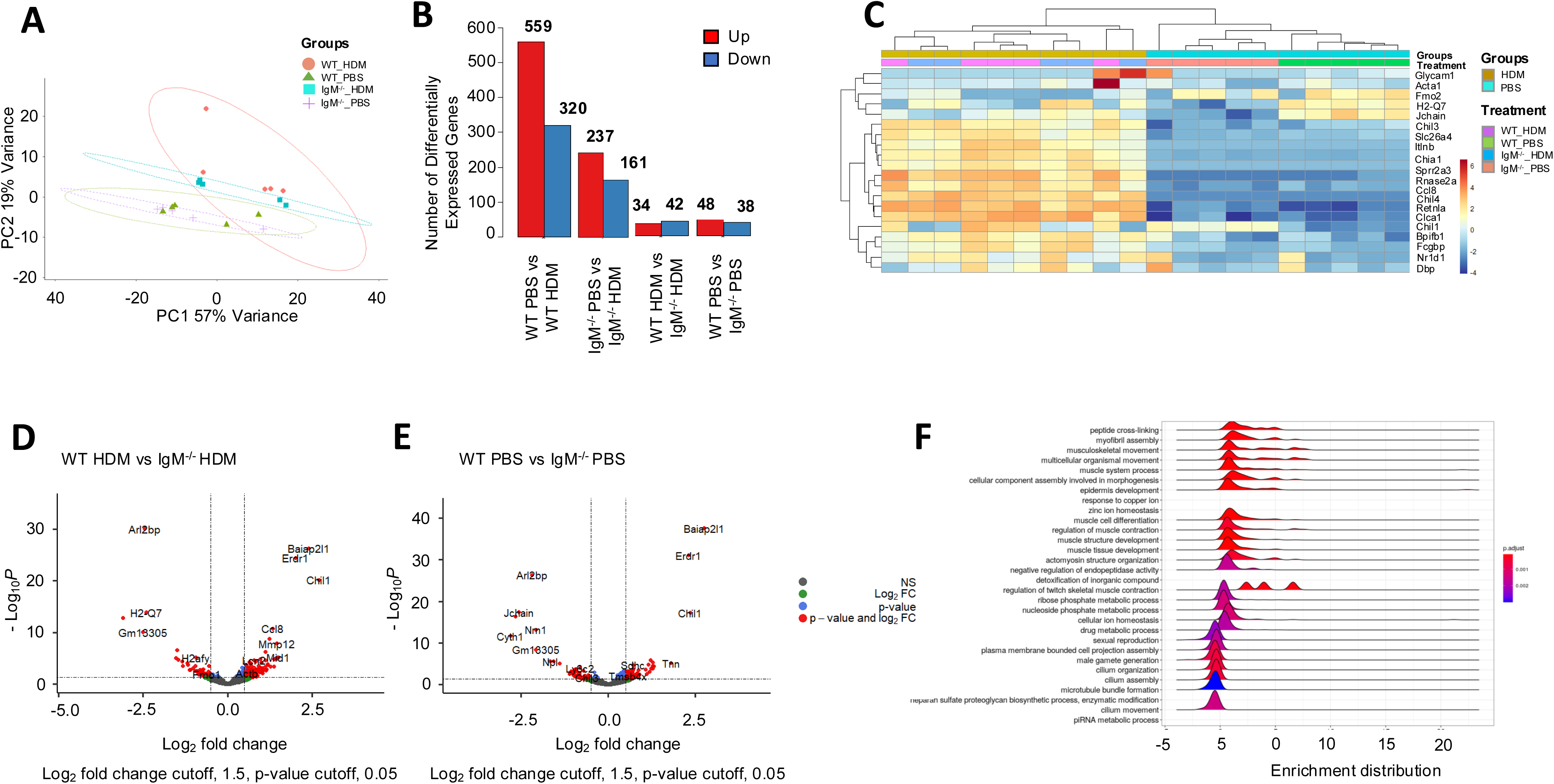
Genes associated with muscle contraction are downregulated in IgM-deficient mice. WT and IgM KO mice were treated as in Figure 1A and RNA was collected from the whole lung for RNA sequencing. **(A)** Principal-component (PC) analysis showing variation in the global gene expression profiles across the different groups. PC1 (60%) and PC2 (18%), which capture the greatest variation in gene expression, are shown. Orange colour represents WT HDM, green colour represents WT PBS, Blue colour represents IgM KO HDM and purple crosses represent IgM KO PBS. Each dot represents an individual mouse. **(B)** Number of differentially expressed genes between groups. **(C)** Heatmaps depicting the differently expressed genes between WT and IgM KO samples from HDM-treated and PBS mice ranked based on hierarchical clustering. **(D**-**E)** Volcano plots: numbers and colour relate to genes that have an adjusted *p* value <0.05. Blue, significantly downregulated; red, significantly up regulated; grey, non-differentially expressed. P values were adjusted for multiple testing using the Benjamini-Hochberg method. **(D)** represent changes between WT and IgM KO treated with HDM and **(E)** represents changes between WT and IgM KO treated with saline. **(F)** Gene set enrichment analysis (GSEA) of hallmark gene sets from the Molecular Signatures Database of the Broad Institute, showing the normalized enrichment scores (NES) for lung RNA-Seq data from WT mice.

### *BAIAP2L1* is expressed in lung cells in contact with airway smooth muscle cells

We decided to focus on *Baiap2l1* also known as Inverse-bin-amphiphysin-Rvs (I-BAR)-domain-containing protein insulin receptor tyrosine kinase substrate (IRTKS), which has been shown to promote actin polymerisation and microvilli length in the intestinal epithelial cells (Postema et al., 2018). We first verified whether BAIAP2L1 was also downregulated at the protein level in IgM-deficient mouse lungs challenged with HDM (Figure 5A). We found lower expression levels of BAIAP2L1 in IgM-deficient mice compared to WT mice challenged with HDM by western blotting, although this was not significant (Figure 5A). Human Protein Atlas search suggested that BAIAP2L1 is expressed by multiple cell types including fibroblasts, skeletal muscle, smooth muscle and macrophages (Abo et al., 2020). We then investigated by immunofluorescence which cell types within the lung could be expressing BAIAP2L1 (Figure 5B). We found BAIAP2L1 to be closely expressed within structural cells in close contact to smooth muscle cells (less than 10μm), a cell type known to be essential in bronchoconstriction. Airway smooth muscle together with the extracellular matrix contributes to smooth muscle hypertrophy which causes the narrowing of the airways. These cells by histology appear to be separated due to fixation methods but are mashed together, especially during the remodelling of the airways (James et al., 2012). IgM deficiency did not impact this expression in cells in close contact with smooth muscle (Figure 5B). Because both western blot and immunofluorescent were inconclusive, we also verified the expression of BAIAP2L1 via flow cytometry (gating in Figure 5-figure supplement 1A) and found it to be expressed mainly by alpha-smooth muscle cells and increased in expression in HDM treated mice compared to PBS control mice (Figure 5C). Furthermore, we showed a reduction in BAIAP2L1 expression amongst alpha-smooth muscle cells but not other cells (CD45 positive or alpha-smooth muscle negative cells, Figure 5-figure supplement 1B) in IgM-deficient mice compared to WT mice treated with HDM, although this did not reach significance (Figure 5C). Overall, this data suggested that BAIAP2L1 is reduced at RNA and protein level and likely influenced the reduction of AHR in IgM-deficient mice. This was consistent with gene set enrichment in our RNA seq data where there was a dominance of genes associated with muscle contraction which regulates AHR.

**Figure 5.**
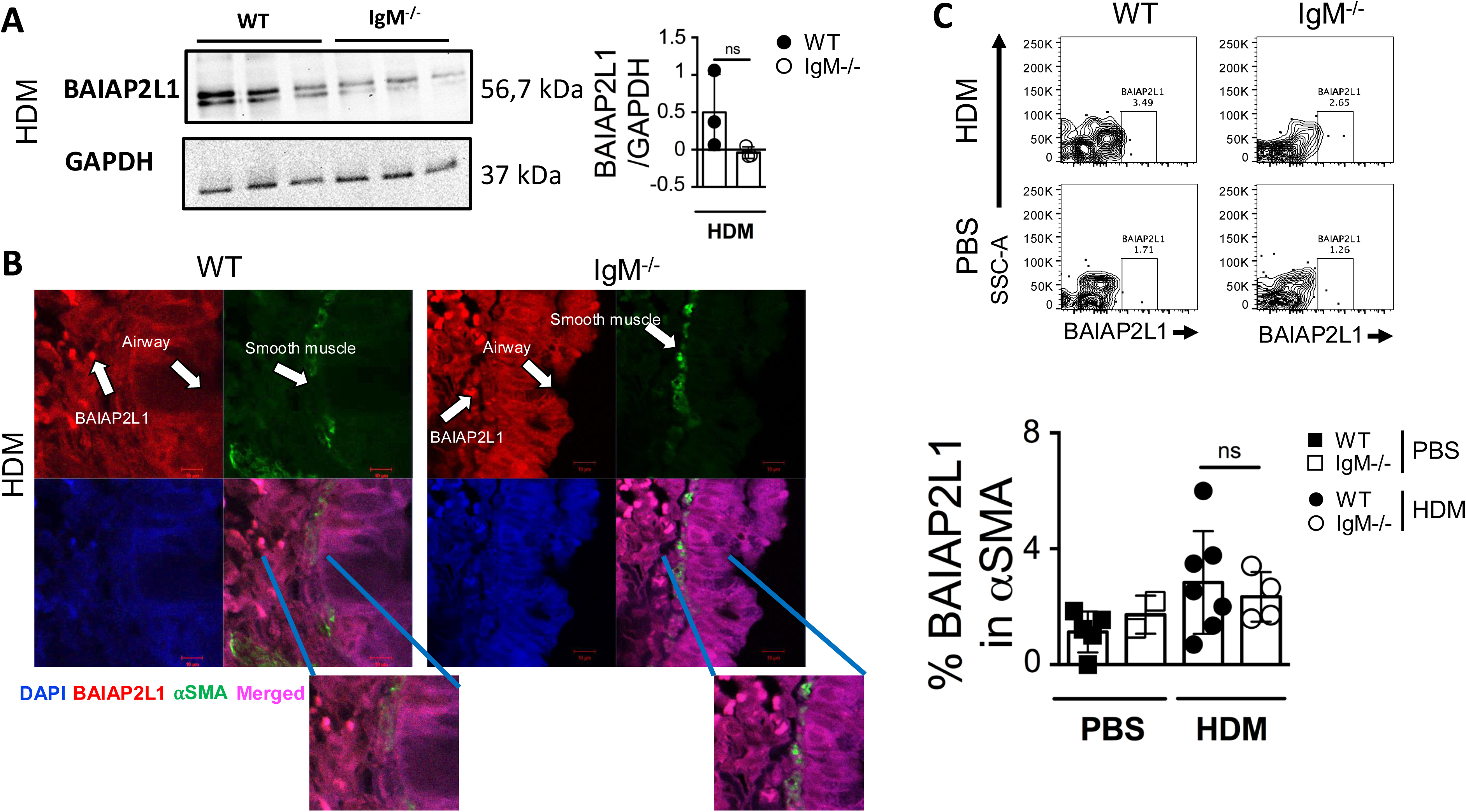
*BAIAP2L1* is expressed in close contact to smooth muscle. **(A)** Mouse lungs were homogenised in RIPA buffer and blotted on nitrocellulose. Rabbit anti-human BAIAP2L1 and mouse anti-GAPDH was used as primary antibody. Lines 1-3 is WT mice, lines 4-6 is IgM KO mice sensitised and challenged with HDM. **(B)** Lung sections from WT and IgM KO mice sensitised and challenged with HDM were immunostained for nuclei stained DAPI (Blue), anti-BAIAP2L1 (red), α smooth muscle actin (green) and merged images (Magenta). Insert below shows zoomed in image of merged BAIAP2L1 and α smooth muscle actin. **(C)** Representative flow cytometry plots showing BAIAP2L1 expression (Live^+^Singlets^+^CD45^-^α SMA^+^BAIAP2L1^+^) in WT and IgM KO treated with HDM or PBS. Quantification of % BAIAP2L1 in α smooth muscle actin is shown. Shown are representative images from 2 independent experiments (n= 3 mice per group).

### CRISPR-Cas9 deletion of *BAIAP2L1* leads to reduced airway smooth muscle contraction at a single cell

To understand how *Baiap2l1*, one of the genes downregulated in IgM-deficient mice, could influence AHR, we resorted to an *in vitro* model that allowed us to measure contraction using fluorescently labelled elastomeric contractible surface (FLECS) technology (Pushkarsky et al., 2018) (Figure 6A). We opted for airway smooth muscle as it has been shown to contribute to the narrowing of the airway during an allergic attack and together with neighbouring cells such as extracellular matrix and immune cells are critical in hypertrophy (James et al., 2012). We chose the human bronchial smooth muscle cell (BSMCs) line as *BAIAP2L1* is expressed in structural cells including muscle and epithelial cells, and actin together with myosin is essential in muscle contraction (Abo et al., 2020). To this end, we used CRISPR-Cas9 technology to knock down *BAIAP2L1* in BSMCs (Figure 6-figure supplement 1A-B). We also included ERDR1 as one of the genes that were also downregulated in IgM-deficient mice. BSMCs (160 000/well) were stimulated with 10 ng/mL of human TNF-α and 100 ng/mL IL-13, followed by transfection with ribonucleoprotein (RNP) complexes containing single guide RNA targeting *BAIAP2L1* and Cas9 for 48h (Figure 6A). Transfected cells were seeded onto elastomeric patterns, stimulated with the same stimulants for 3h and fixed in 4% paraformaldehyde, followed by staining with DAPI and Phalloidin before imaging on Stellarvision microscope and quantification (Figure 6A). We validated the deletion of *BAIAP2L1* by PCR and showed reduced expression of *BAIAP2L1* in acetylcholine (ACh) and TNF-α stimulated cells transfected with *BAIAP2L1* sgRNA when compared to scramble sgRNA (Figure 6 - figure supplement 1C, line 3 and 4 compared to line 5 and 6). We also detected low expression of BAIAP2L1 in unstimulated sgRNA scramble and sgRNA *BAIAP2L1* transfected cells (Figure 6-figure supplement 1C, lines 1 and 2). Transfections with sgRNA did not impact cell viability (Figure 6-figure supplement 1D). As expected, BSMCs displayed a higher level of contraction in TNF-α or IL-13 stimulated cells compared to unstimulated cells (Figure 6B-D). In BSMCs transfected with sgRNA- *BAIAP2L1* and -*ERDR1*, we saw a significant reduction in contraction levels in TNF-α stimulated cells when compared to scramble sgRNA transfected and stimulated cells (Figure 6B, C). We saw no differences in contraction between scramble sgRNA and sgRNA-*BAIAP2L1* and -*ERDR1* stimulated with IL-13 (Figure 6D).

**Figure 6.**
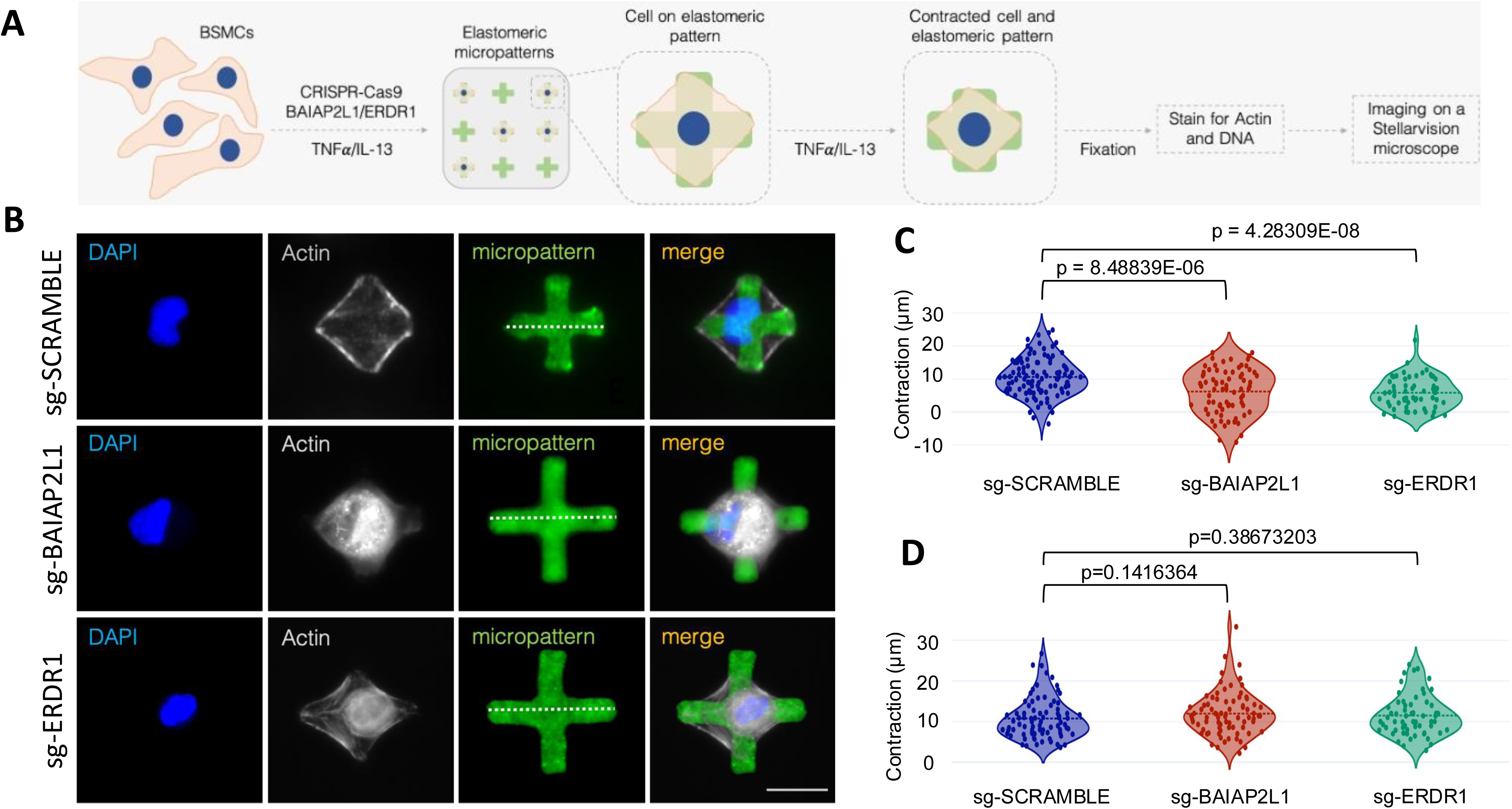
CRISPR-based deletion of *BAIAP2L1* leads to reduced smooth muscle contraction at a single-cell level. **(A)** Bronchial smooth muscle cells (1.6×10^5^ cells/well) were transfected with CRISPR-Cas9 single guide RNAs (scramble, *BAIAP2L1* and *ERDR1*), stimulated with recombinant human IL-13 (100 ng/mL) and TNF-α (10 ng/mL) for 48 h. Cells were then transferred to elastomeric micropatterns, stimulated again with rIL-13 and rTNF-α and fixed before imaging on a StellarVision microscope. **(B)** Representative images of single BSMCs on micropatterns from scramble, *BAIAP2L1* and *ERDR1* stimulated with 10 ng/mL TNF-α. DNA was stained with DAPI, actin fibers with Phaloidin-565 and elastomeric micropatterns are coated in Fibronectin-488. Merged images are shown on the right. **(C)** Violin plots showing contraction of 50-100 cells/condition stimulated with 10 ng/mL TNF-α, individual dots represent a single cell contraction, where blue is scramble sgRNA, red is *BAIAP2L1* sgRNA and green is *ERDR1* is sgRNA. **(D)** Violin plots showing contraction of 50-100 cells/condition stimulated with 100 ng/mL IL-13, individual dots represent a single cell contraction, where blue is scramble sgRNA, red is *BAIAP2L1* sgRNA and green is *ERDR1* is sgRNA. Shown is mean ±SEMs from two pooled experiment (n=50 - 100). Significant differences between groups were performed by student t-test (Mann-Whitney) and *p* value is shown.

### *BAIAP2L1* deletion but not *ERDR1* is essential in acetylcholine-induced smooth muscle contraction

Contraction of the ASM is central to AHR as it controls airway diameter and airflow which can increase resistance (Lambrecht and Hammad, 2014). Airway smooth muscle contractility is induced when extracellular factors such as ACh bind through muscarinic 3 acetylcholine receptor (m_3_AChR), activating a signalling cascade leading to calcium accumulation and contraction (Ouedraogo and Roux, 2014). We transfected BSMCs with sgRNA-*BAIAP2L1*, -*ERDR1* or scrambled and stimulated cells with 10 µM ACh (Figure 7A). We saw a significant reduction in BSMC contraction transfected with sgRNA-*BAIAP2L1* when compared to scramble sgRNA transfected and ACh stimulated (Figure 7B-C). We saw no significant differences in BSMC contraction between sgRNA-*ERDR1* transfected and scramble sgRNA transfected and ACh stimulated cells (Figure 7B-C). Taken together, this data indicates that *BAIAP2L1* a gene that was downregulated in IgM-deficient mice has a stimulant-specific role in inducing airway contraction of bronchial smooth muscle cells during asthma.

**Figure 7.**
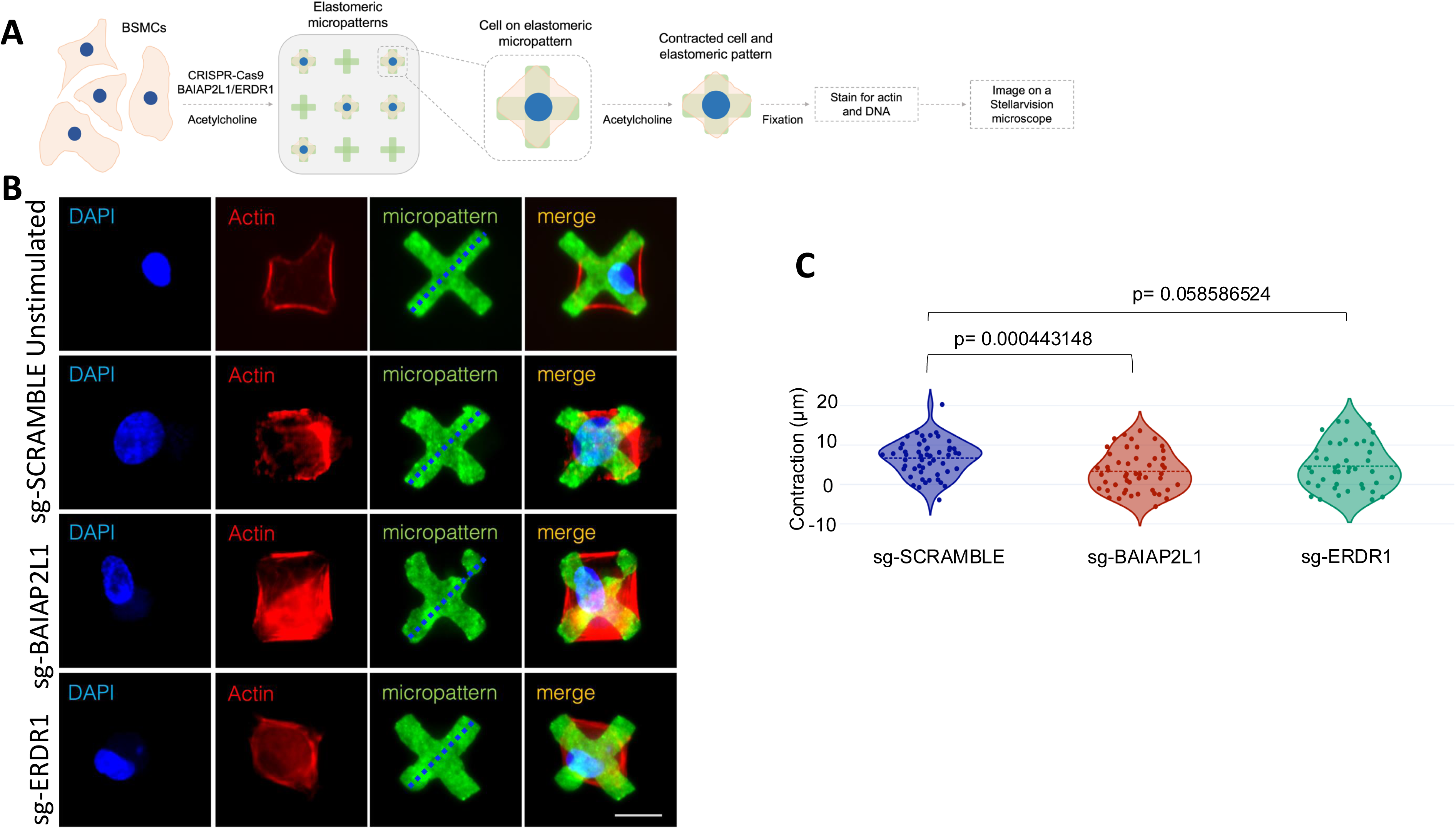
CRISPR-based deletion of *BAIAP2L1* reduces smooth muscle contraction upon stimulation with acetylcholine. **(A)** Bronchial smooth muscle cells (1.6×10^5^ cells/well) were transfected with CRISPR-Cas9 single guide RNAs (scramble, *BAIAP2L1* and *ERDR1*), stimulated with Acetylcholine (10µM) for 48 h. Cells were then transferred to elastomeric micropatterns, stimulated again with ACh (10µM) and fixed before imaging on a StellarVision microscope. **(B)** Representative images of single BSMCs on a single micropattern from unstimulated, scramble, *BAIAP2L1* and *ERDR1* stimulated with ACh (10µM). Shown are DAPI, actin (red), a green fluorescent micropattern and merged images. **(C)** Violin plots showing contraction of 50-100 cells/condition stimulated with ACh (10µM) individual dots represent a single cell contraction, where blue is scramble sgRNA, red is BAIAP2L1 sgRNA and green is ERDR1 is sgRNA. Shown is mean ±SD from 1 representative experiment of 2 independent experiments (n=50 - 100). Significant differences between groups were performed by student t-test (Mann-Whitney) and *p* value is shown.

## Discussion

Here, we described an unexpected function of IgM in the regulation of bronchoconstriction. We show by RNA sequencing that in the lung tissue of IgM-deficient mice, there is a reduction in genes (*Baiap2l1* and *Erdr1*) associated with actin cytoskeleton and re-arrangement, a key factor in smooth muscle contraction and AHR. We used single-cell force cytometry and CRISPR-Cas9 technology to validate these genes in the human BSM cell line and show as a proof of concept that deletion of *BAIAP2L1* reduced muscle contraction in a stimulant-specific manner.

B cells play a complex role in allergic asthma, earlier studies using B cell-deficient mice showed a redundant role of B cells in OVA-induced allergic airway inflammation and airway hyperreactivity (Hamelmann et al., 1999; Korsgren et al., 1997; MacLean et al., 1999). We and others have recently shown using a more complex allergen HDM relevant to human asthma that antigen load is crucial in B cell function (Dullaers et al., 2017; Habener et al., 2021; Hadebe et al., 2021). We explored these possibilities in the context of IgM deficiency and observed a profound reduction in AHR. This was in complete contrast to what has been observed in B cell-deficient mice and in IgE or FcεRI-deficient mice (Dullaers et al., 2017; Habener et al., 2021; McKnight et al., 2017). We also found a similar reduction in AHR at higher doses of HDM, albeit less pronounced. Reduction in AHR was not only specific to HDM, but we also found reduced AHR in IgM-deficient mice sensitised with OVA complexed to alum and challenged with OVA, although less pronounced. AHR reduction in IgM-deficient mice was also seen in the acute papain model which only activates innate responses including innate lymphoid cells (Darby et al., 2021). Interestingly, this reduction in AHR was not associated with reduced allergic airway inflammation including eosinophilia, mucus production or Th2 cells in all models tested, which was unexpected considering these features are key in AHR. B cell-deficient mice show a reduction in eosinophilia and antigen-specific Th2 cells at low doses of HDM, which leads to reduced AHR (Dullaers et al., 2017). Similar findings are observed in mice lacking IL-4Rα specifically in B cells when challenged with HDM (Hadebe et al., 2021).

IgM deficiency did not impact B cell development in primary and secondary lymphoid tissues and accumulation of follicular and germinal centre B cells. Interestingly, there was a lack of class switching to IgG1 and IgE isotypes despite increased expression of IgD. The lack of class switching to IgG1 and IgE contrasts with earlier reports, which suggested that IgD can largely replace IgM for class switching to other isotypes, resulting in delayed neutralising IgG1 against vesicular stomatitis virus (VSV) (Lutz et al., 1998; Ochsenbein et al., 1999). To get a better understanding of which IgM function was responsible for AHR, we showed that the transfer of naïve wild-type mice serum to IgM-deficient mice could restore IgE production and HDM-specific IgE and IgG1. This is consistent with what has been observed in the context of influenza viral infections, where the transfer of purified or serum IgM into B cell-deficient mice restored IgM-induced viral neutralisation (Jayasekera et al., 2007). We believe that naïve sera transferred to IgM-deficient mice were able to bind to the surface of B cells via IgM receptors (FcμR / Fcα/μR), which are still present on IgM-deficient B cells, and this signalling is sufficient to facilitate Class Switch Recombination (CSR). This was also confirmed by the ability of B cells to reach germinal centres in IgM-deficient mice during HDM exposure. Our IgM KO mouse lacks both membrane-bound and secreted IgM and transferred serum contains at least secreted IgM which can bind to B cell surfaces. Of course, we can’t rule out that transferred sera from WT mice also contains some IgG1 which can facilitate class switching to IgE when transferred to IgM-deficient mice. Despite the ability of wild-type serum to restore IgE, it was not enough to restore AHR, which may be due to the difficulties in restoring IgM to normal levels (200-800 µg). Our findings on the redundant role of IgE were consistent with previous studies where IgE nor its high-affinity receptor (FcεRI) were essential in AHR in an HDM or OVA model (McKnight et al., 2017). The lack of functional role of serum-transferred IgE was consistent with earlier findings on *H. polygyrus* transfer of immune serum where IgE was found not to be essential in protection against *H. polygyrus* re-infection (Wojciechowski et al., 2009). To resolve other endogenous factors that could have potentially influenced reduced AHR in IgM-deficient mice, we resorted to busulfan chemical irradiation to deplete bone marrow cells in IgM-deficient mice and replace bone marrow with WT bone marrow. While it is well accepted that busulfan chemical irradiation partially depletes bone marrow cells, in our case it was not possible to pursue other irradiation methods due to changes in ethical regulations and that fact that mice are slow to recover after gamma rays irradiation. Busulfan chemical irradiation allowed us to show that we could mostly restore AHR in IgM-deficient recipient mice that received donor WT bone marrow when challenged with low dose HDM.

B cell isotype IgD has been shown to promote Th2 cells and OVA-, papain- and NP-specific humoral responses specifically IgG1 and IgE through basophil activation (Shan et al., 2018). IgD has also been shown to inhibit IgE-mediated basophil activation through downregulation of genes associated with cytoskeleton organisation such as signal transducers phosphoinositide-3 kinase, RAS, and RHO (Shan et al., 2018). Our RNA sequencing data showed downregulation of *Baiap2l1* and *Erdr1* in IgM-deficient mice, genes which have been shown to associate with actin cytoskeleton and re-arrangement, (Abo et al., 2020; Houh et al., 2016; Postema et al., 2018). Contraction of the ASM is central to AHR as it controls airway diameter and airflow which can increase resistance (Perkins et al., 2011). ASM activated by cytokines such as IL-13 or TNF-α or bronchoconstrictor ACh induces cytosolic calcium accumulation (Moulton and Fryer, 2011; Perkins et al., 2011). Ca^2+^ release results in a cascade of events that involves kinases and phosphorylation of key molecules such as actin and myosin involved in smooth muscle contraction (Ouedraogo and Roux, 2014). Gene set enrichment analysis suggested that genes downregulated in IgM-deficient mice were involved in processes such as muscle contraction, muscle twitching, skeletal muscle movement. As a proof of concept we validated the involvement of *BAIAP2L1* in bronchial smooth muscle contraction using a high throughput method that allows us to measure contraction of 1000s of single cells (Pushkarsky et al., 2018). When we deleted *BAIAP2L1* using CRISPR-Cas9, we showed that deletion of *BAIAP2L1* in BSMCs reduced smooth muscle contraction when stimulated with TNF-α and ACh, but not IL-13. Deletion of ERDR1 had minor impact on muscle contraction across different stimulants suggesting a stimulant and gene-specific role. IRTKS/BAIAP2L1 has been shown to regulate microvilli elongation by forming a complex with actin-regulating protein EPS8 (Postema et al., 2018). In this setting IRTKS elongate microvilli via distinct mechanisms that require functional WH2 and SH3 domains, which binds EPS8, an F-actin capping and bundling protein (Postema et al., 2018). We have not fully dissected how *BAIAP2L1* could be regulated by IgM, but it is clear that through its actions to regulate actin bundling, it is involved in muscle contraction which requires actin and myosin head interaction (summarised in Figure 8).

**Figure 8.**
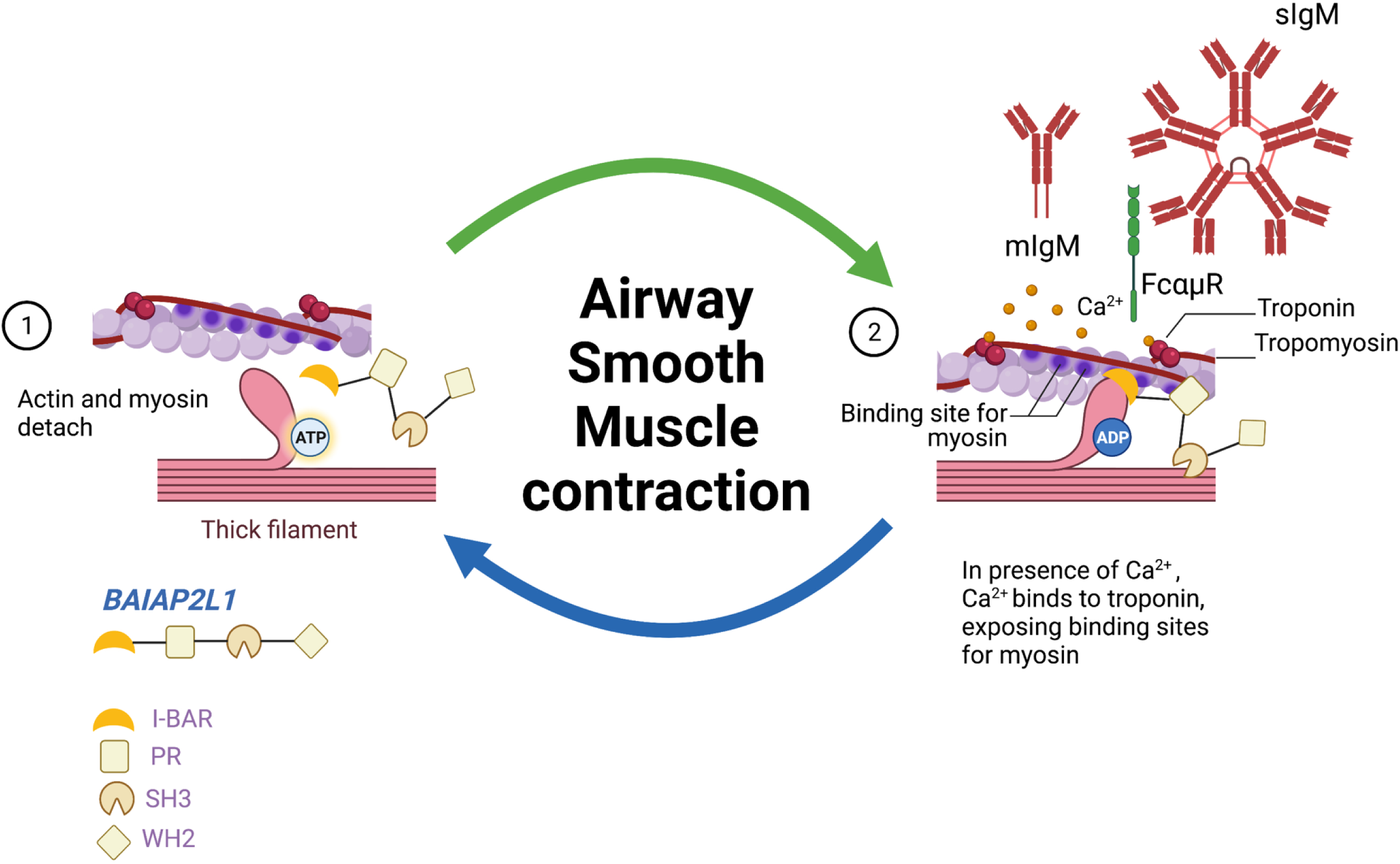
Working model showing how *Baiap2l1* could influence muscle contraction through influencing myosin and actin filament interaction. Image adapted from Biorender.

We speculate that IgM can directly activate smooth muscle cells by binding a number of its surface receptors including FcμR, Fcα/μR and pIgR (Liu et al., 2019; Nguyen et al., 2017b; Shibuya et al., 2000). IgM binds to FcμR strictly, but shares Fcα/μR and pIgR with IgA (Liu et al., 2019; Michaud et al., 2020; Nguyen et al., 2017b). Both Fcα/μR and pIgR can be expressed by non-structural cells at mucosal sites (Kim et al., 2014; Liu et al., 2019). We would not rule out that the mechanisms of muscle contraction might be through one of these IgM receptors, especially the ones expressed on smooth muscle cells(Kim et al., 2014; Liu et al., 2019). Certainly, our future studies will be directed towards characterizing the mechanism by which IgM potentially activates the smooth muscle.

We believe these findings report for the first time an independent function of IgM from its natural class switching and non-specific binding to microbial components. We believe that IgM function in regulating ASM may be indirect through other unknown factors but does not involve microbiota as treatment of mice with a mixture of antibiotics did not restore AHR. Early vaccination with bacterial species, such as group *A streptococcus* containing GlcNAc or β-1,3-glucans can protect adult mice against *A. fumigatus*-induced allergic asthma (Kin et al., 2012). This is mainly through conserved germline-encoded IgM antibodies, which have broad specificities to common allergens containing GlcNAc moieties such as Dermatophytes (Kearney et al., 2015; New et al., 2020). Together these findings demonstrate for the first time an important function of IgM in regulating airway hyperresponsiveness independent of the presence of T helper 2 allergic inflammation. These findings have implications for future treatment of allergic asthma through bronchodilators.

## Materials and Methods

### Mice

IgM-deficient homozygotes mice on Balb/C background and on C57BL/6 background were originally described in (Lutz et al., 1998) and were backcrossed at least 10 generations in house at the University of Cape Town (Lutz et al., 1998). Wild type on Balb/C (The Jackson Laboratory, RRID:IMSR_JAX:000651) and C57BL/6 (The Jackson Laboratory, RRID:IMSR_JAX:000664) backgrounds were used as a littermate control. Congenic wild type Balb/C mice (CByJ.SJL(B6)-Ptprca/J, RRID:IMSR_JAX:006584) were backcrossed to at least 10 generations at the University of Cape Town. B cell-deficient mice (muMt B6.129S2-Ighmtm1Cgn/J, RRID:IMSR_JAX:002288) was originally described in (Kitamura et al., 1991) and was backcrossed at least 10 generations to a Balb/C background at the University of Cape Town. Mice were housed in independently ventilated cages under specific pathogen-free conditions at the University of Cape Town Animal Facility. All mice were used at eight to 10 weeks of age and animal procedures were performed according to the strict recommendation by the South African Veterinary Council and were approved by the University of Cape Town Animal Ethics Committee (Reference number 014/019, 018/013 and 022/014).

### House Dust-mite induced allergic airway disease

A high dose and a low dose HDM treatment schedule were used to induce symptoms of allergic asthma in mice (Hadebe et al., 2021). Mice were anaesthetised with ketamine (Anaket-V; Centaur Labs, Johannesburg, South Africa) and xylazine (Rompun; Bayer, Isando, South Africa). For low-dose HDM, mice were sensitised intratracheally (i.t.) on day 0 with 1 µg of HDM (Stellergens Greer Laboratories, Lenoir, U.S.A.) and intranasally challenged with 3 ug HDM on days 8, 9, 10, 11 and 12. For high-dose HDM, mice were with HDM 100 ug and challenged with HDM 10 ug. AHR was measured on day 15. After the procedure, mice were euthanised and tissue samples were collected for analysis.

### Adoptive transfer of naïve B cells

Spleens were collected from naïve congenic CD45.1 Balb/C mice and passed through 40μm strainer to obtain single-cell suspensions. Cells were stained with FITC-B220 and APC-CD19 for 30min at 4°C. A dead cell exclusion dye (7AAD) was added before sorting on BD FACS Aria I to at least 96% purity. 2-5 x 10^6^ cells were adoptively transferred intravenously (i.v.) into IgM KO recipient mice a day before HDM sensitisation.

### Adoptive transfer of naïve serum

Naïve wild-type mice were euthanised and blood was collected via cardiac puncture before being spun down (5500rpm, 10min, RT) to collect serum. Serum (200μL) was injected intraperitoneally into IgM-deficient mice. Serum was injected intraperitoneally at day −1, 0, and a day before the challenge with HDM (day 10).

### Busulfan Bone marrow chimeras

WT (CD45.2) and IgM KO (CD45.2) congenic mice were treated with 25 mg/kg busulfan (Sigma-Aldrich, Aston Manor, South Africa) per day for 3 consecutive days (75 mg/kg in total) dissolved in 10% DMSO and Phosphate buffered saline (0.2mL, intraperitoneally) to ablate bone marrow cells. Twenty-four hours after last administration of busulfan, mice were injected intravenously with fresh bone marrow (10×10^6^ cells, 100μL) isolated from hind leg femurs of either WT (CD45.1) or IgM KO mice (Montecino-Rodriguez and Dorshkind, 2020). Animals were then allowed to complement their haematopoietic cells for 8 weeks. In some experiments the level of bone marrow ablation was assessed 4 days post-busulfan treatment in mice that did not receive donor cells. At the end of experiment level of complemented cells were also assessed in WT and IgM KO mice that received WT (CD45.1) bone marrow.

### Ovalbumin-induced allergic airway inflammation

Mice were sensitised intraperitoneally with (50µg in 200µl) of ovalbumin (OVA) adsorbed to 0.65% alum (Sigma-Aldrich, Aston Manor, South Africa) on days 0, 7, 14. On days 23, 24, 25, mice were intranasally challenged with 100µg of OVA under anaesthesia with ketamine (Anaket-V; Centaur Labs, Johannesburg, South Africa) and xylazine (Rompun; Bayer, Isando, South Africa). AHR was measured on day 26. After the procedure, mice were euthanized with halothane and tissue samples collected for analysis.

### Papain-induced lung inflammation

Mice were anaesthetized with isoflourane (3L/min) briefly before being challenged with 50μL of 25 μg of Papain (Sigma-Aldrich, Aston Manor, South Africa) on days 1, 2 and 3 or PBS. AHR was measured on day 4. After the procedure, mice were euthanized with halothane and tissue samples collected for analysis.

### Airway Hyperresponsiveness

Airway resistance and elastance of the whole respiratory system (airways, lung chest wall) after intranasal challenge was determined by forced oscillation measurements as described previously (Kirstein et al., 2015) with the Flexivent system (SCIREQ, Montreal, Canada) by using the single compartment (‘‘snapshot’’) perturbation. Measurements were carried out with increasing doses of acetyl-β-methylcholine (methacholine, Sigma-Aldrich, Aston Manor, South Africa) (0, 5, 10, 20 and 40 mg/mL) for Balb/C or (0, 20, 40, 80 160 and 320 mg/mL) for C57BL/6. Differences in the dose-response curves were analysed by repeated-measures Two-way ANOVA with the Bonferroni post-test. Only mice with acceptable measurements for all doses (coefficient of determination >0.90) were included in the analysis.

### Flow cytometry

Single-cell suspensions were prepared from lymph nodes in Roswell Park Memorial Institute (RPMI) media (Gibco, Paisley, United Kingdom) by passing them through 100µm strainer. To obtain single cell suspensions from lung tissues, a left lobe was digested for 1 hour at 37°C in RPMI containing 13 mg/mL DNase I (Roche, Randburg, South Africa) and 50 U/mL collagenase IV (Gibco, Waltham, Massachusetts) and passed through 70µm strainer. Antibodies used in these experiments included, phycoerythrobilin (PE)- conjugated anti-Siglec-F (clone, E50-2440), anti-IL-5 (clone, TRFK5), anti-CD44 (clone, KM114), anti-T and B cell activation antigen (clone, GL7), anti-CD43 (clone, S7), FITC-conjugated anti-Ly6G (clone, 1A8), anti-IgD (clone, 11-26C2a), IL-4 (clone, 11B11), anti-PD-1-(clone, 29F.1A12), PerCP Cy5.5-conjugated anti-Ly6C (clone, AL-21), -CD45.1 (clone, A20), anti-IL-17 (clone, TC11-18H10), anti-mouse alpha muscle actin (Abcam, ab8211-500), Allophycocyanin (APC)- conjugated anti-CD11c (clone, HL3), anti-CD5 (clone, 53-7.3), BV421 conjugated anti-CD11b (clone, M1/70), anti-CD62L (clone, MEL-14), anti-IgG1 (clone, A110-1), AlexaFlour 700- conjugated anti-CD3ε (clone, 145-2C11) -anti-IFN-γ (clone, XMG1.2), BV510-anti-CD4 (clone, RM4-5) and anti-B220 (clone, RA3-6B2), APC-Cy7-conjugated anti-CD19 (clone, 1D3) and anti-CD8 (clone, 53-6.7), BV786 conjugated anti-IgE (clone, R35-72) and anti-IL-33R (ST2) (clone, U29-93), biotin-conjugated anti-IgM (clone, AF-78), anti-CD95 (clone, Jo2), anti-CD249 (clone, BP-1), anti-CD45 (clone, 30-F11) were purchased from BD Pharmingen (San Diego, CA). PE-Cynanine7 anti-F4/80 (clone, BM8), anti-IL-13 (clone, eBio13A), anti-CXCR5 (clone, L138D7), AlexaFlouro 700- conjugated anti-MHC II (clone, M5/114), Rabbit anti-BAIAP2L1 (Abcam, PA554000), Live/dead Fixable Yellow stain (Qdot605 dead cell exclusion dye) were purchased from eBiosciences. Biotin-labelled antibodies were detected by Texas Red conjugated PE (BD Biosciences). PE-Goat anti-Rabbit IgG (Abcam, ab72465) was used to detect Rabbit anti-BAIAP2L1. For staining, cells (1 x 10^6^) were stained and washed in PBS, 3% FCS FACS buffer. For intracellular cytokine staining, cells were restimulated with phorbal myristate acetate (Sigma-Aldrich) (50 ng/mL), ionomycin (Sigma-Aldrich) (250ng/mL), and monensin (Sigma-Aldrich) (200mM in IMDM/10% FCS) for 5h at 37°C then fixed in 2% PFA, permeabilised with Foxp3 transcriptional factor staining buffer kit (eBioscience) before intracellular staining with appropriate cytokine antibodies and acquisition through LSR Fortessa machine (BD Immunocytometry system, San Jose, CA, USA) and data was analysed using Flowjo software (Treestar, Ashland, OR, USA).

### Histology

Left upper lung lobes was fixed in 4% formaldehyde/PBS and embedded in paraffin. Tissue sections were stained with periodic acid-Schiff for mucus secretion, and haematoxylin and eosin (H&E) stain for inflammation. Slides were scanned at 20x magnification on the virtual slide VS120 microscope (Olympus, Japan). Downstream processing of images was done through Image J (FIJI) for image extraction at series 15 and Ilastik software was used for mucus area quantification on whole lung sections. The data shown are representative of 1 experiment of 3 independent experiments (n = 5-7 mice per experiment).

### Antibody and cytokine ELISAs

Antibody ELISAs were carried out as previously described (Kirstein et al., 2015) using 10 μg/ml HDM to coat for specific IgGs. Total IgE in serum was measured using anti-mouse IgE (BD Biosciences, 553413) to coat, mouse IgE (κ, anti-TNP, BD Biosciences, 557079) as standard and biotin anti-mouse IgE (BD Biosciences, 553419) as a secondary antibody.

For *in vitro* cytokine production analysis, single-cell suspensions were prepared from mediastinal lymph nodes of HDM-treated and littermate control mice. Cells (2×10^5^ cells, in 200µL) were incubated for 5 days in RPMI/10% FCS (Delta Bioproducts, Kempton Park, South Africa) in 96-well plates. Cells were either stimulated with HDM (30µg/mL) or plate bound anti-CD3 (10µg/mL) and supernatants were collected after a 5-day incubation period. Concentrations of IL-4, IL-5 (BD Biosciences) and IL-13 (R&D Systems, Minneapolis, Minn), were measured using ELISA assays according to the manufacturer’s protocol.

### RNA Extraction

Small lung lobe was frozen in Qiazol (Qiagen, Germany) and stored at −80°C. Total RNA was isolated from the lysate using miRNeasy Mini kit (Qiagen, Germany) according to the manufacturer’s instructions. RNA quantity and purity were measured using the ND-1000 NanoDrop spectrophotometer (ThermoScientific, DE, USA).

### cDNA Synthesis and RT-qPCR

For *Muc5a* gene expression analysis, 100 ng total RNA was reverse transcribed into cDNA using Transcriptor First Strand cDNA Synthesis Kit (Roche, Germany) according to the manufacturer’s instructions. Quantitative real-time PCR (RT-qPCR) was performed using LightCycler® 480 SYBR Green I Master (Roche, Germany) and *Muc5a* primers (IDT, CA, USA). Fold change in gene expression was calculated by the ΔΔCt method and normalized to *Actb* which was used as an internal control.

### Whole lung RNA sequencing

Whole lung RNA was extracted using RNAeasy kit (Qiagen, Germany) according to the manufacturer’s instructions. We used Agilent 2100 Bio analyzer (Agilent RNA 6000 Nano Kit) to do the total RNA sample QC: RNA concentration, RIN value,28S/18S and the fragment length distribution. The first step in the workflow involved purifying the poly-A containing mRNA molecules using poly-T oligo attached magnetic beads. Following purification, the mRNA was fragmented into small pieces using divalent cations under elevated temperature. The cleaved RNA fragments were copied into first strand cDNA using reverse transcriptase and random primers. This was followed by second strand cDNA synthesis using DNA Polymerase I and RNase H. These cDNA fragments had addition of a single ‘A’ base and subsequent ligation of the adapter. The products were then purified and enriched with PCR amplification. PCR yields were quantified by Qubit and pooled samples together to make a single strand DNA circle (ssDNA circle), which gave the final library. DNA nanoballs (DNBs) were generated with the ssDNA circle by rolling circle replication (RCR) to enlarge the fluorescent signals at the sequencing process. The DNBs were loaded into the patterned nanoarrays and pair-end reads of 100 bp were read through on the DNBseq platform for the following data analysis study. For this step, the DNBseq platform combines the DNA nanoball-based nanoarrays and stepwise sequencing using Combinational Probe-Anchor Synthesis Sequencing Method.

### Bioinformatics workflow

The ribosomal RNA (rRNA) was first removed using SortMeRNA. We then did the Fastq file quality control using Fastqc and multiqc software to assess the quality of the raw reads followed by adapter trimming using Trim galore. The reads were then aligned to the mouse reference genome (mm10_UCSC_20180903) using STAR aligner. The map read counts was then extracted using featurecounts. The gene differential analysis was conducted using the DEseq2. The genes with LFC >= 2 and adjusted p-value <= 0.05 was used to do the gene ontology over-representation analysis done using clusterProfiler (Wu et al., 2021) Bioconductor package. The Benjamini-Hochberg method was used for multiple test correction.

### Human bronchial Airway Smooth Muscle cell culture

A healthy human bronchial smooth muscle cell line (BSMC) was obtained from Lonza (Catalog #: CC-2576) and cultured in Smooth Muscle Growth Medium (Lonza) supplemented with Insulin (CC-4021D), human Fibroblastic Growth Factor-B (CC-4068D), Gentamicin sulfate-Amphotericin-1000 (CC-4081D), 5% Foetal Bovine Serum (CC-4102D), human Epidermal Growth Factor (CC-4230D) (all purchased from Lonza) in a T25 flask (Lonza) until confluence. After two passages in the T75 flask, confluent cells were seeded at 1.6×10^5^ BSMC cells onto 24-well trays and immediately transfected with CRISPR/Cas9 single guide RNAs. No mycoplasma contamination was detected during the time of culturing. We verified human BSMCs by staining with alpha smooth muscle antibody.

### CRISPR/Cas9 single guide RNA transfections

single guide RNAs targeting human *BAIAP2L1* (# CD.Cas9.CXVQ6494.AA); *ERDR1*, (# CD.Cas9.YFVV2490.AA); HPRT Negative Control (#Alt-R® CRISPR-Cas9crRNA) were purchased (IDT, CA, USA via WhiteScientific PTY LTD). Single guide RNA (1 μM) and Cas9 enzyme (1 μM) in Opti-MEM medium (Life Technologies™, Carlsbad, CA, USA) were transfected using Lipofectamine RNAi Max 1000 (Thermo Scientific) into BSMC cells (1.6 ×10^5^) per well and either stimulated with Acetylcholine (10μM), recombinant human IL-13 (100 ng/mL), recombinant human TNF-α (10 ng/mL) or left unstimulated for 48 hours at 37°C, 5% CO_2_ incubator. Gene deletion was confirmed by DNA extraction (Wizard Genomic DNA Purif. Kit, Promega) and PCR amplification of target genes using *BAIAP2L1* forward (GTCCCGGGGGCCCGA) and reverse (AAGCGCCCAAGAATGTGGGG) primers, and product run on a 1.2% agarose gel.

### Cytotoxic detection Assay

To measure cytotoxicity after single guide RNA transfection, lactate dehydrogenase (LDH) was measured in supernatants collected at 48 hours post transfection using cytotoxicity detection kit ^PLUS^ Assay (CYTODET-RO, Roche) according to manufacturer’s instructions and plates were read at 490 nm.

### Single-cell force cytometry using Fluorescently Labelled Elastomeric Contractible Surface (FLECS) technology

BSMCs were seeded on Fibronectin-coated elastomeric micropatterns (Forcyte Biotechnologies #F2AX0G03Y, 50 microns, Alexa Fluor 488 - bound Fibrinogen, 24-well plate) at a concentration of 75,000 cells per well in Smooth muscle growth medium (SmGM) (Lonza) and left to adhere and spread for 90 minutes. Unattached cells were then removed and fresh SmGM medium supplemented with rhTNF-α (10 ng/mL)/rhIL-13 (100 ng/mL)/ACh (10μM) was added. BSMCs were stimulated for 3 hrs before being fixed in pre-warmed to 37°C 4% PFA. Fixed samples were washed and then stained with ATTO-565 phalloidin (ATTO-TEC) and DAPI (Life Technologies) and imaged on a StellarVision microscope using Synthetic Aperture Optics technology (Optical Biosystems). All images were analyzed using FiJi software by measuring the length of each micropattern per condition after stimulation and subtracting the length of the unstimulated micropattern of the same condition.

### Immunofluorescent

Left upper lung lobes were fixed in 4% formaldehyde/PBS and embedded in paraffin. Tissue sections (5 μm) were deparaffinised and hydrated in water before antigen retrieval in boiling 10 mM Citrate buffer (pH 6, pressure cooker for 2 mins). Tissue was incubated with Rabbit anti-BAIAP2L1 (Abcam, PA554000) in PBS-Tween (1:250, overnight, 4°C) and FITC anti-mouse alpha muscle actin (Abcam, ab8211-500, 1:100, overnight, 4°C). Tissues were washed with PBS-Tween before incubation with secondary antibody PE-Goat anti-Rabbit IgG (Abcam, ab72465) in PBS-Tween (1:500, RT in the dark for 30min). Tissues were washed with PBS-Tween before mounting in DAPI mounting media (Thermofisher). Fluorescent images were captured at 60x magnification using LSM880 Airy Scan (Zeiss, UK).

### Western blotting

Protein harvest: RIPA lysis buffer supplemented with protease inhibitors was added to the lung tissue. The whole-cell lysates were incubated on ice for 20 min and vortexed every 5 min for 60 seconds. Then, the lysate was centrifuged at 3000 rpm for 15 minutes at 4°C, and the supernatant was collected. Total protein concentrations were determined using the Pierce BCA Assay kit.

SDS-PAGE: 20 µg of each protein sample was mixed with Laemmli buffer and heated for 10 minutes at 100 °C. Equal amounts of protein samples were loaded into 5% acrylamide stacking gel and a 12% acrylamide separating gel along with 5 µL PageRulerTM Plus Prestained protein ladder (catalogue #: 26619, Thermo ScientificTM) and separated by electrophoresis at 100 V. Proteins were transferred onto nitrocellulose membranes (Bio-Rad) using the Bio-Rad Mini Trans-Blot® Cell according to the manufacturer’s instructions.

After protein transfer, the nitrocellulose membranes were stained with Ponceau stain for approximately 1-3 minutes to visualize if the proteins had been transferred. Once the protein bands were identified, the membrane was washed with ddH_2_O until 80% of the Ponceau staining is removed. The remaining 20% of the stain allowed the accurate segmentation of the membrane to separate high molecular weight proteins from low molecular weight proteins. The nitrocellulose membranes were blocked with 5 % (w/v) non-fat milk, 0.01% Tween-20 in TBST) on the shaker at room temperature for one hour. The low and high molecular weight membranes proteins were incubated with respective primary antibodies (Rabbit anti-human BAIAP2L1, PA5-54000, Invitrogen) or (Goat anti-mouse GAPDH, sc 365062, Santa Cruz) overnight shaking in the 4°C cold room. Membranes were washed 4 times with 1xTBST in 15-minute intervals. Horseradish peroxidase (HRP) secondary antibodies were added to their respective membranes and incubated for one hour at room temperature, with shaking. subsequently, membranes were washed 3 times in 1xTBST in 15 minutes intervals. To visualise the protein bands on autoradiographic film (Santa Cruz Biotechnology®), Lumino Glo® substrate kit was added to each membrane. Briefly, the substrate luminol was oxidised by hydrogen peroxide in the presence of the catalyst HRP to yield a chemiluminescent product.

### Plating of faecal pellets to confirm a reduction in bacterial colonies following antibiotic treatment

The faecal pellet was collected from each individual mouse treated with antibiotics as well as control groups not treated with antibiotics. The faecal pellet was weighed and resuspended in sterile PBS at a concentration of 50 mg/mL. The sample was resuspended until homogenous, and 1/100 dilution was made in sterile PBS. 50μL of 1/100 dilutions was plated on tryptic soy agar plates and spread evenly across the plate before incubating at 37°C for 16 hrs. Individual colonies were counted manually.

### Statistical analysis

P-values were calculated in GraphPad Prism 6 (GraphPad Software, Inc) by using nonparametric Mann-Whitney Student’s t-test or Two-way ANOVA with Bonferroni’s post-test for multiple comparisons, and results are presented as standard error of the mean (SEM) or mean of standard deviation (SD). Differences were considered significant if P was <0.05.

## Acknowledgements

We thank the UCT Research Animal Facility for maintaining mice. Wendy Green and Munadia Ansari for genotyping mice. We are grateful to Lizette Fick, Matt Darby and Raygaana Jacobs for their excellent histology services and the UCT Confocal Microscopy Core Facilities. We are grateful to Ronnie Dreyer for the excellent cell sorting and Flow Cytometry Core Facility. We thank FlowJo for proving free service to Africa.

## Data availability statement

The data related to this paper can be found at NCBI Sequence Read Archive (SRA) number <u>PRJNA1288527</u>, Single cell force cytometry data can be found at Figshare under DOI 10.6084/m9.figshare.29832320; 10.6084/m9.figshare.29614157; 10.6084/m9.figshare.29832263.

---

**Figure 1 -figure supplement 1.**
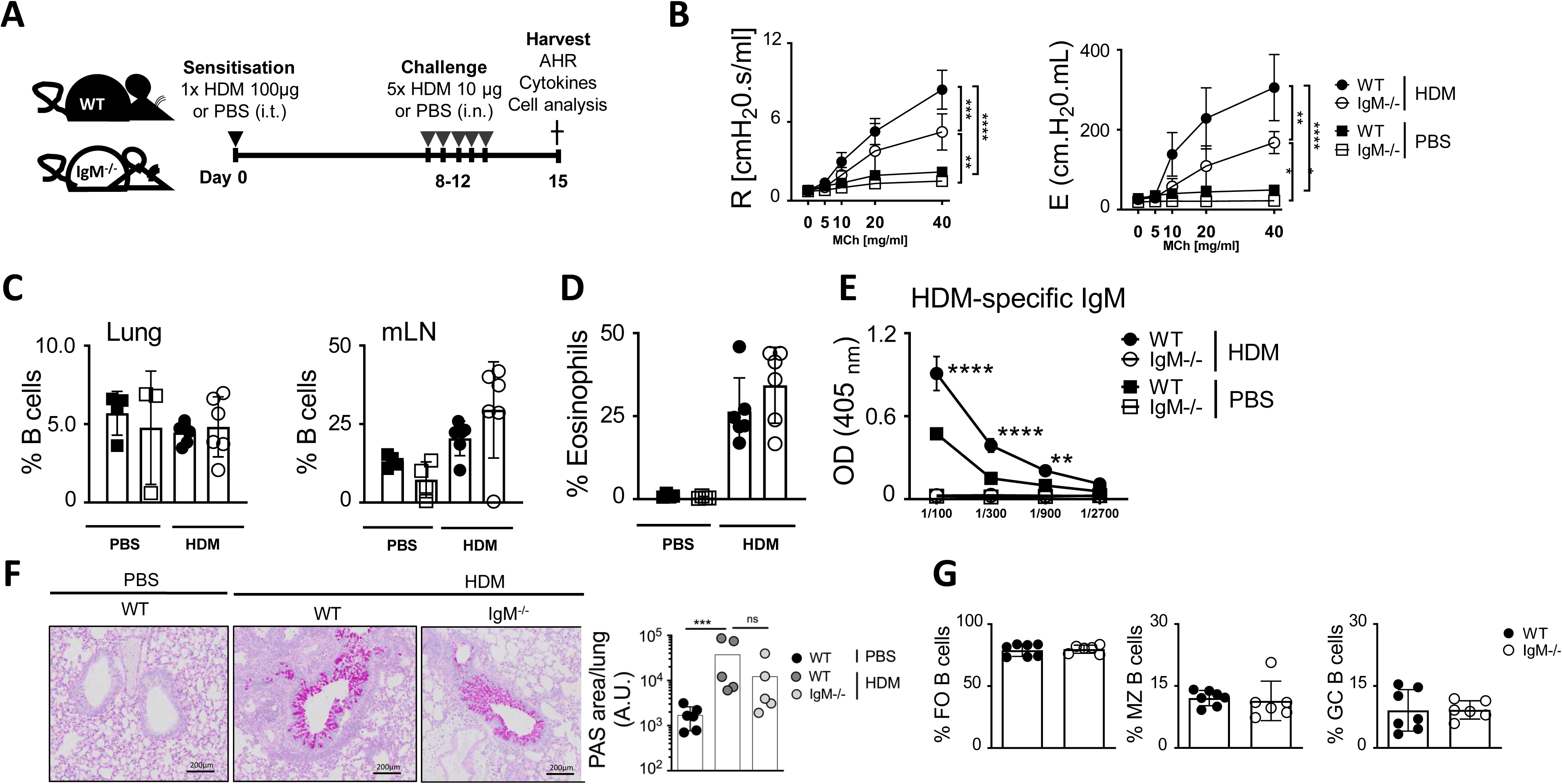
IgM deficiency leads to reduced airway hyperresponsiveness, but eosinophils and B cells are unaffected. **(A)** Schematic diagram showing sensitisation and challenge protocol where mice (IgM KO) and wild-type littermate control (WT) were sensitised with HDM 100 μg intratracheally on days 0 and challenged with HDM 10 μg on days 8-12. Analysis was done on day 15. **(B)** Airway resistance and elastance were measured with increasing doses of acetyl methacholine (0 −40 mg/mL). **(C)** Frequencies of lung and mediastinal lymph node B cells (live^+^B220^+^CD19^+^MHCII^+^) and were stained and analysed by Flow cytometry. **(D)** Frequencies of lung eosinophils (live^+^Siglec-F^+^CD11c^-^) were stained and analysed by Flow cytometry. **(E)** HDM-specific IgM in serum. **(F)** Histology analyses of lung sections (scale 200μm), stained with periodic acid-Schiff. A.U., Arbitrary units **(G)** Quantification of follicular B cells, Marginal Zone B cells and Germinal Centre B cells. Shown is mean ±SEMs from two pooled experiment (n= 10-12). Significant differences between groups were performed by Two way ANOVA with Benforroni post-test and are described as: *p<0.05, **p<0.01, ***p<0.001, ****p< 0.0001.

**Figure 1 -figure supplement 2.**
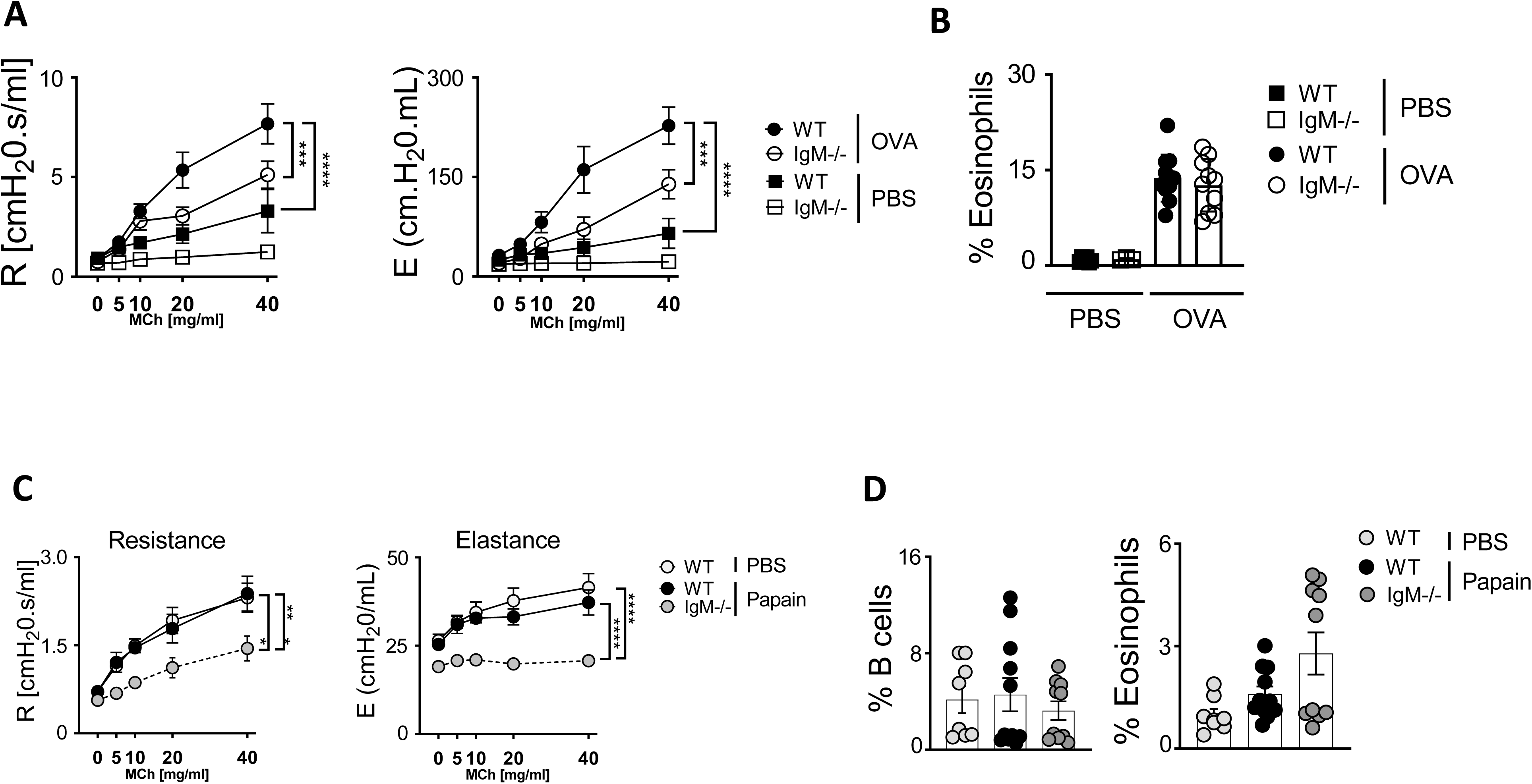
Reduced airway hyperresponsiveness in lgM-deficient mice is independent of allergen. IgM KO and wild type (WT) were sensitised intraperitoneally with (50µg in 200µl) of ovalbumin (OVA) adsorbed to 0.65% alum on days 0, 7, 14. On days 23, 24, 25, mice were intranasally challenged with 100µg of OVA. AHR was measured on day 26. **(A)** Airway resistance and elastance were measured with increasing doses of acetyl methacholine (0 −40 mg/mL). **(B)** Frequencies of lung eosinophils (live^+^Siglec-F^+^CD11c^-^) were stained and analysed by Flow cytometry. **(C)** IgM KO and wild type (WT) were challenged with 25 μg of papain on days 1, 2 and 3 or PBS. AHR was measured on day 4. Airway resistance and elastance were measured with increasing doses of acetyl methacholine (0 −40 mg/mL). **(D)** Frequencies of lung B cells (live^+^B220^+^CD19^+^MHCII^+^) and eosinophils (live^+^Siglec-F^+^CD11c^-^) were stained and analysed by Flow cytometry. Shown is mean ±SEMs from two pooled experiment (n= 10-12). Significant differences between groups were performed by Two way ANOVA with Benforroni post-test and are described as: **p<0.01, ***p<0.001, ****p< 0.0001.

**Figure 1 -figure supplement 3.**
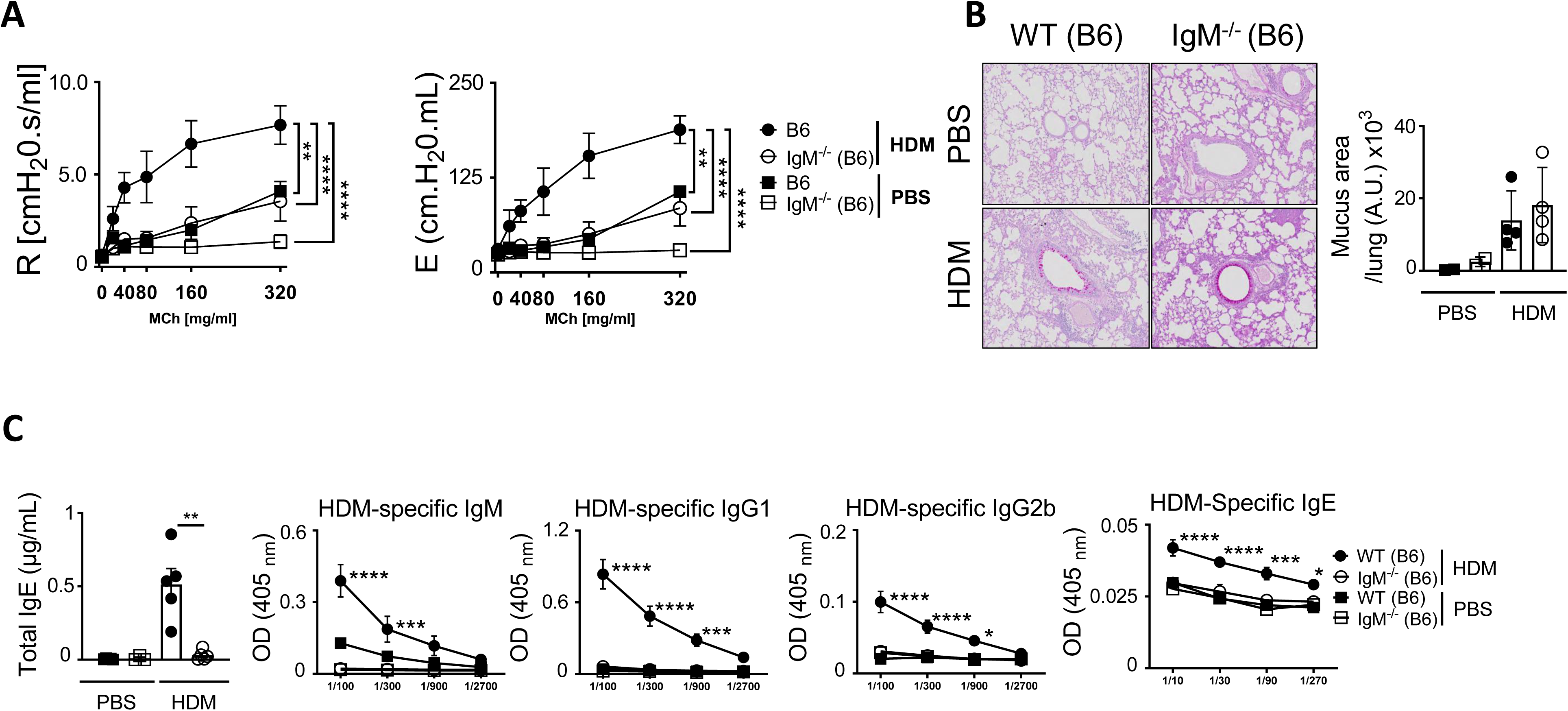
Reduced airway hyperresponsiveness in lgM-deficient mice is independent of mouse background in HDM-induced asthma. **(A)** Mice (IgM KO and WT) were sensitised and challenged as in Figure 1, A and Airway resistance and elastance were measured with increasing doses of acetyl methacholine (0 −320 mg/mL). **(B)** Histology analyses of lung sections (magnification x20), stained with periodic acid-Schiff. A.U., Arbitrary units. **(C)** Total IgE, HDM-specific IgM, HDM-specific IgG1, HDM-specific IgG2a and HDM-specific IgE in serum. Shown is mean ±SD from 1 representative experiment of 2 independent experiments (n= 4-6). Significant differences between groups were performed by student t-test (Mann-Whitney) (C) or Two way ANOVA with Benforroni post-test and are described as: *p<0.05, **p<0.01, ***p<0.001, ****p< 0.0001.

**Figure 1 -figure supplement 4.**
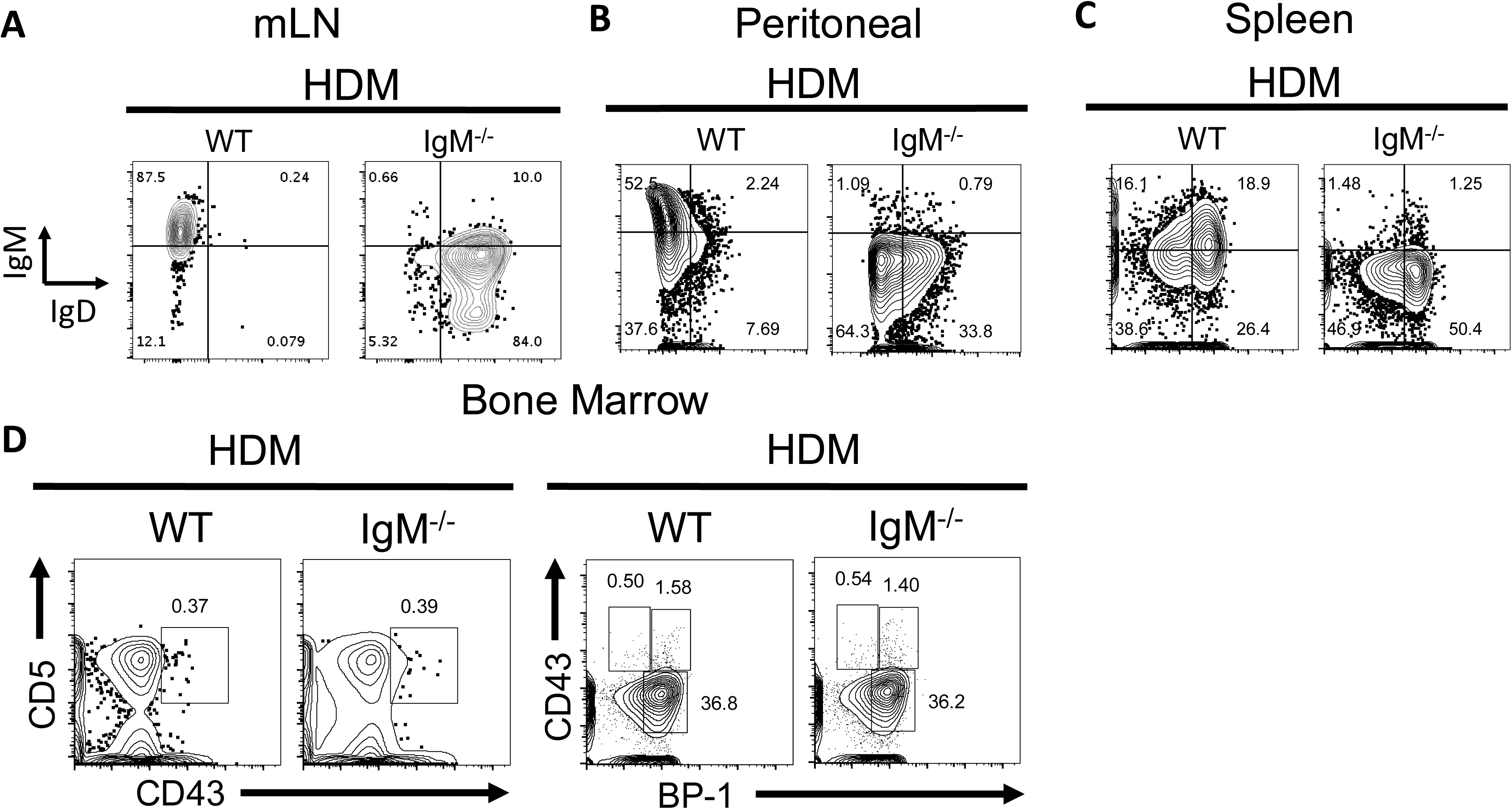
B cell development is not impacted by lack of lgM, however, IgD expression is upregulated in all tissues. **(A)** B cell expression of IgM and IgD in IgM KO and WT mice treated with HDM in mediastinal lymph nodes, (**B)** Peritoneal and (**C)** Spleen. **(D)** Bone marrow was flushed out IgM KO and WT mice treated with HDM and stained for B1 cells (CD5^+^CD43^+^) and pre-B cells (CD43^+^BP-1^+^).

**Figure 2 -figure supplement 1.**
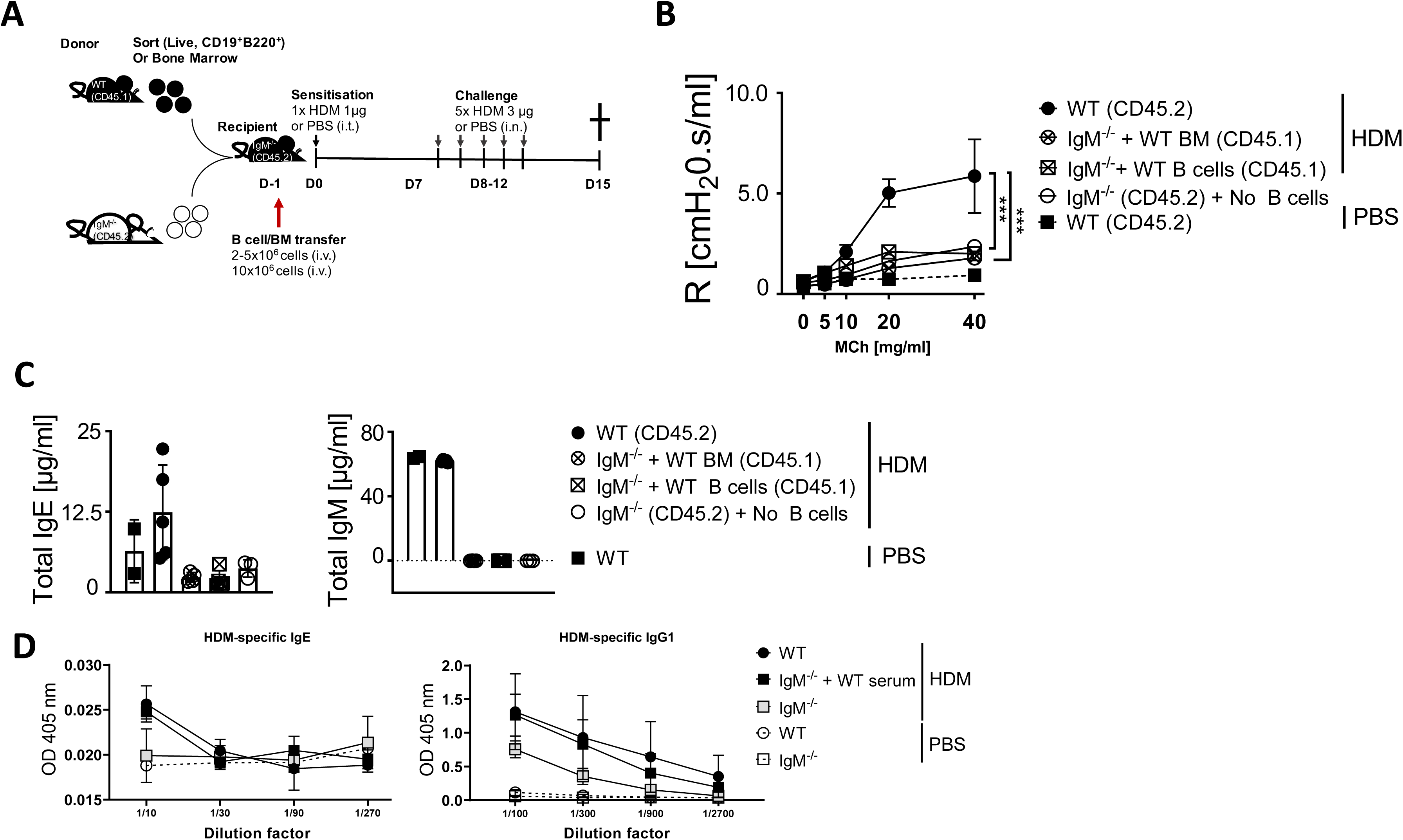
B cell and bone marrow reconstitution do not restore airway hyperresponsiveness in lgM-deficient mice in HDM-induced allergic asthma. **(A)** Schematic diagram showing sorted B cells (live^+^B220^+^CD19^+^) (2-5×10^6^) or bone marrow cells (10×10^6^) transferred from congenic WT (CD45.1) to IgM KO a day before being sensitised as shown in Fig. 1,A. **(B)** Airway resistance was measured with increasing doses of acetyl methacholine (0-40 mg/mL). **(C)** Total IgE and IgM in serum of mice reconstituted with WT B cells or bone marrow. **(D)** House Dust Mite specific IgE and IgG1 in IgM KO mice transferred with WT serum (mice treated as in Figure 2D). Shown is mean ±SD from 1 experiment (n= 4-6). Significant differences between groups were performed by Two-way ANOVA with Benforroni post-test and are described as: ***p<0.001.

**Figure 3 -figure supplement 1.**
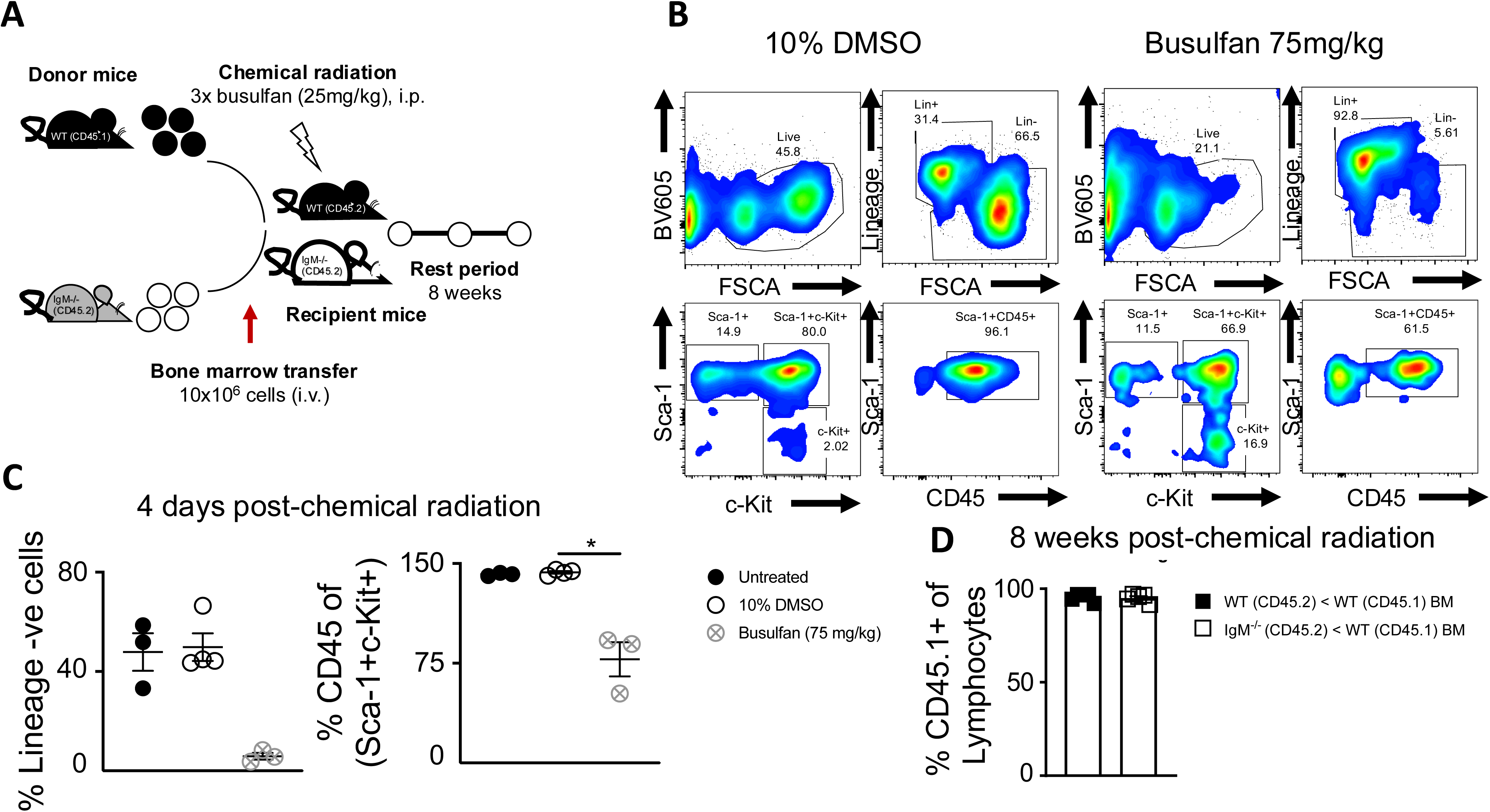
Bone marrow chimaera generation using busulfan to restore IgM function. **(A)** Schematic diagram showing WT and IgM-deficient mice being chemically irradiated with busulfan (25 mg per day for 3 days) and adoptively transferred with congenic WT or syngeneic IgM KO bone marrow (10×10^6^ per mouse intravenously) 24 hours post-last busulfan treatment. Recipient mice (WT to IgM KO) were then rested for 8 weeks before being sensitised as shown in Fig 1a. **(B)** FACS plots of bone marrow cells from either vehicle (10% DMSO) or busulfan (75 mg/kg) treated mice, showing live cells (BV605 live/dead dye), Lineage positive (Lin^+^) and negative cells (Lin^-^), frequencies of Sca-1^+^c-Kit^+^ cells within Lin^+^ and frequencies of CD45^+^ cells within Sca-1^+^c-Kit^+^ cells. **(C)** Quantification of frequencies of lineage negative cells (Lin^-^) and frequencies of CD45^+^ cells within Sca-1^+^c-Kit^+^ cells. **(D)** Quantification of frequencies of CD45.1^+^ cells within live lymphocytes in lung cells of WT or IgM KO busulfan treated recipient mice that received donor WT CD45.1 bone marrow, 8 weeks post busulfan treatment. Shown is the mean ± SD from one experiments (n=5 – 6 per group). Significant differences between groups were performed by Student *t*-test (Mann-Whitney) (C) and are described as: *p<0.05.

**Figure 3 -figure supplement 2.**
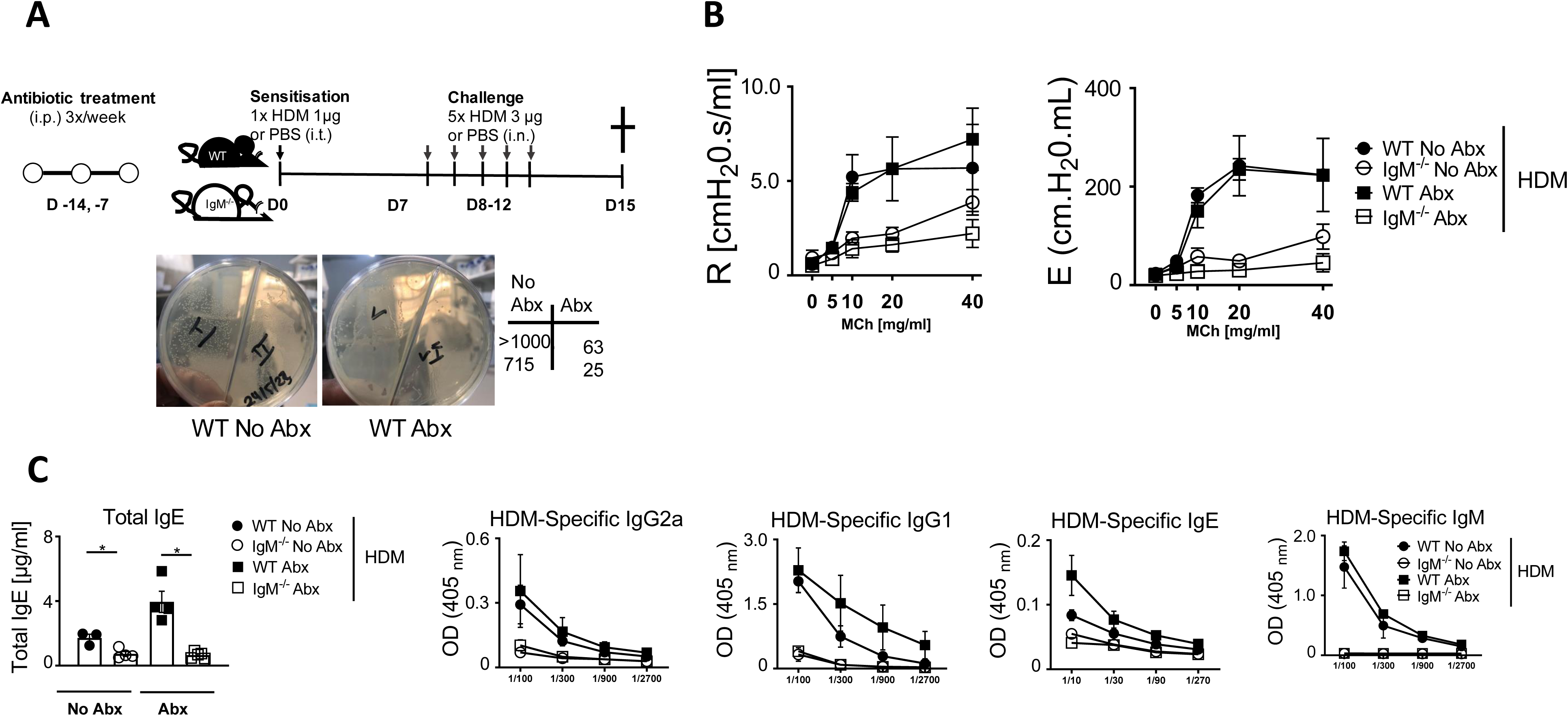
Reduced airway hyperresponsiveness in lgM-deficient mice is independent of microbial influence in HDM-induced asthma. **(A)** Schematic diagram showing treatment of IgM KO and WT mice with antibiotic mixture 3 times a week for 2 weeks via oral gavage. These mice were then sensitised and challenged as in Figure 1A. Depicted are the number of colonies from TSA agar plates grown for 16 hrs at 37°C from faecal samples of WT mice treated (Abx) or not treated with an antibiotic cocktail (No Abx). **(B)** Airway resistance and elastance were measured with increasing doses of acetyl methacholine (0 −40 mg/mL). **(C)** Total IgE, HDM-specific IgG2a, HDM-specific IgG1, HDM-specific IgE and HDM-specific IgM in serum. Shown is mean ±SDs from 1 experiment (n= 4-6). Significant differences between groups were performed by student t-test (Mann-Whitney) and are described as: *p<0.05, **p<0.01, ***p<0.001, ****p< 0.0001. Abx, Antibiotic.

**Figure 4 -figure supplement 1.**
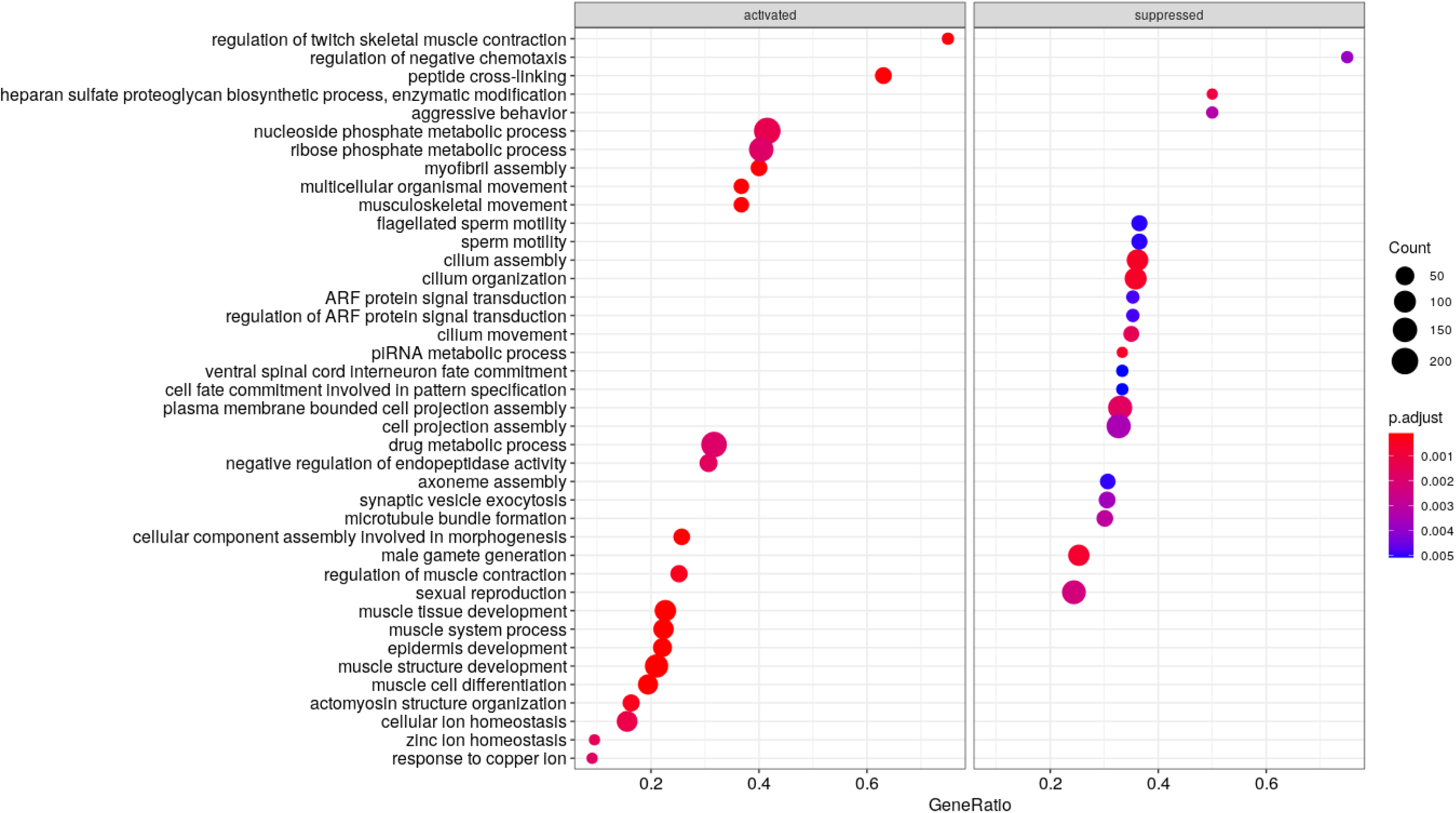
Genes associated with muscle contraction are downregulated in IgM-deficient mice. Gene set enrichment analysis (GSEA) showing gene ratio of activated and suppressed pathways from lung RNA-seq data from WT mice and IgM KO mice. Related to Figure 4, F.

**Figure 5 -figure supplement 1.**
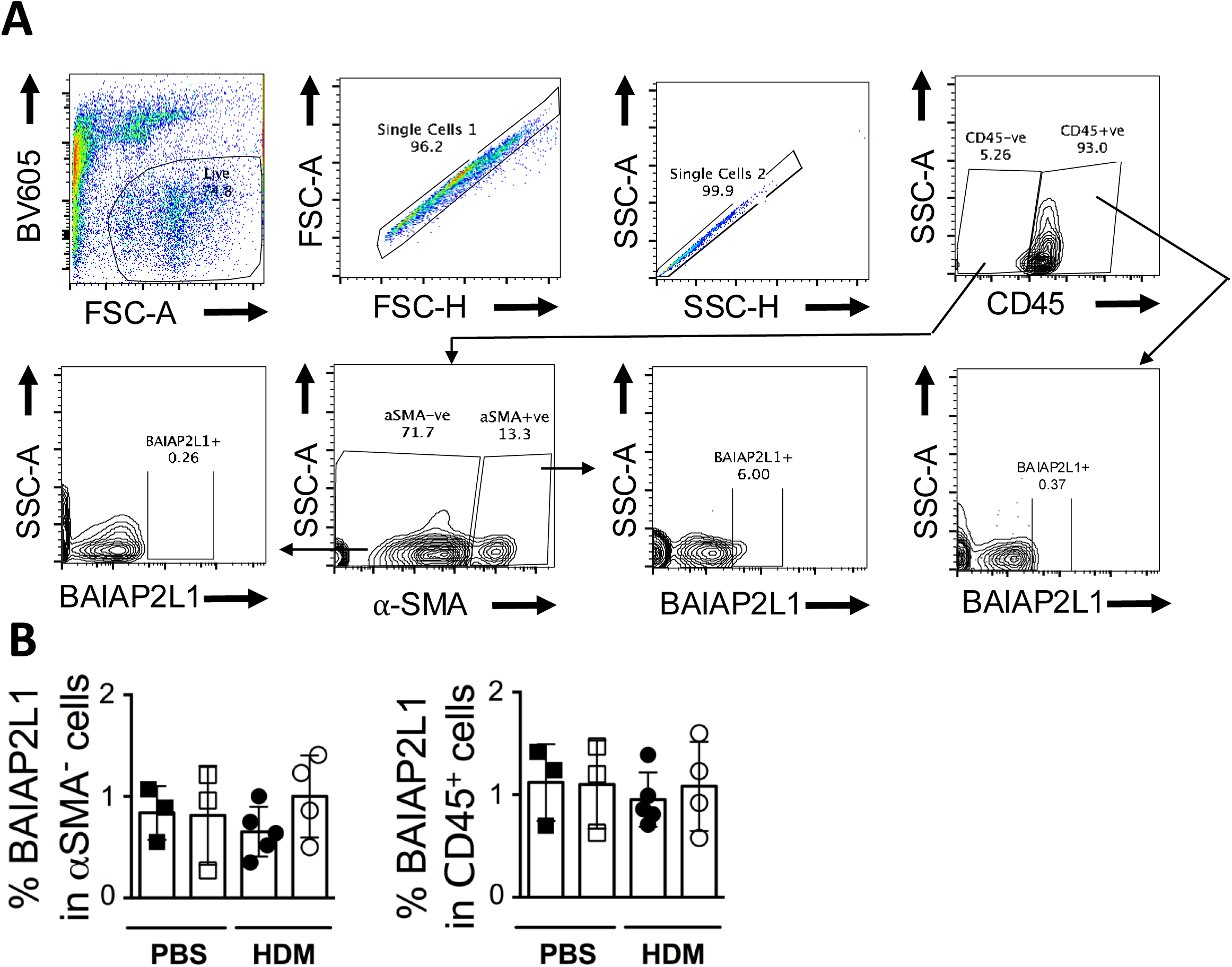
BAIAP2L1 is mainly expressed by alpha-smooth muscle cells. **(A)** Flow cytometry gating strategy showing expression of BAIAP2L1 in CD45+ cells (live^+^Singlets1^+^Singlets2^+^CD45^+^BAIAP2L1^+^), alpha-smooth muscle positive cells (live^+^Singlets1^+^Singlets2^+^CD45^-^αSMA^+^BAIAP2L1^+^) and alpha-smooth muscle negative cells (live^+^Singlets1^+^Singlets2^+^CD45^-^αSMA^-^BAIAP2L1^+^). **(B)** Quantification of expression of BAIAP2L1 in CD45+ cells (live^+^Singlets1^+^Singlets2^+^CD45^+^BAIAP2L1^+^) and alpha-smooth muscle negative cells (live^+^Singlets1^+^Singlets2^+^CD45^-^αSMA^-^BAIAP2L1^+^).

**Figure 6 -figure supplement 1.**
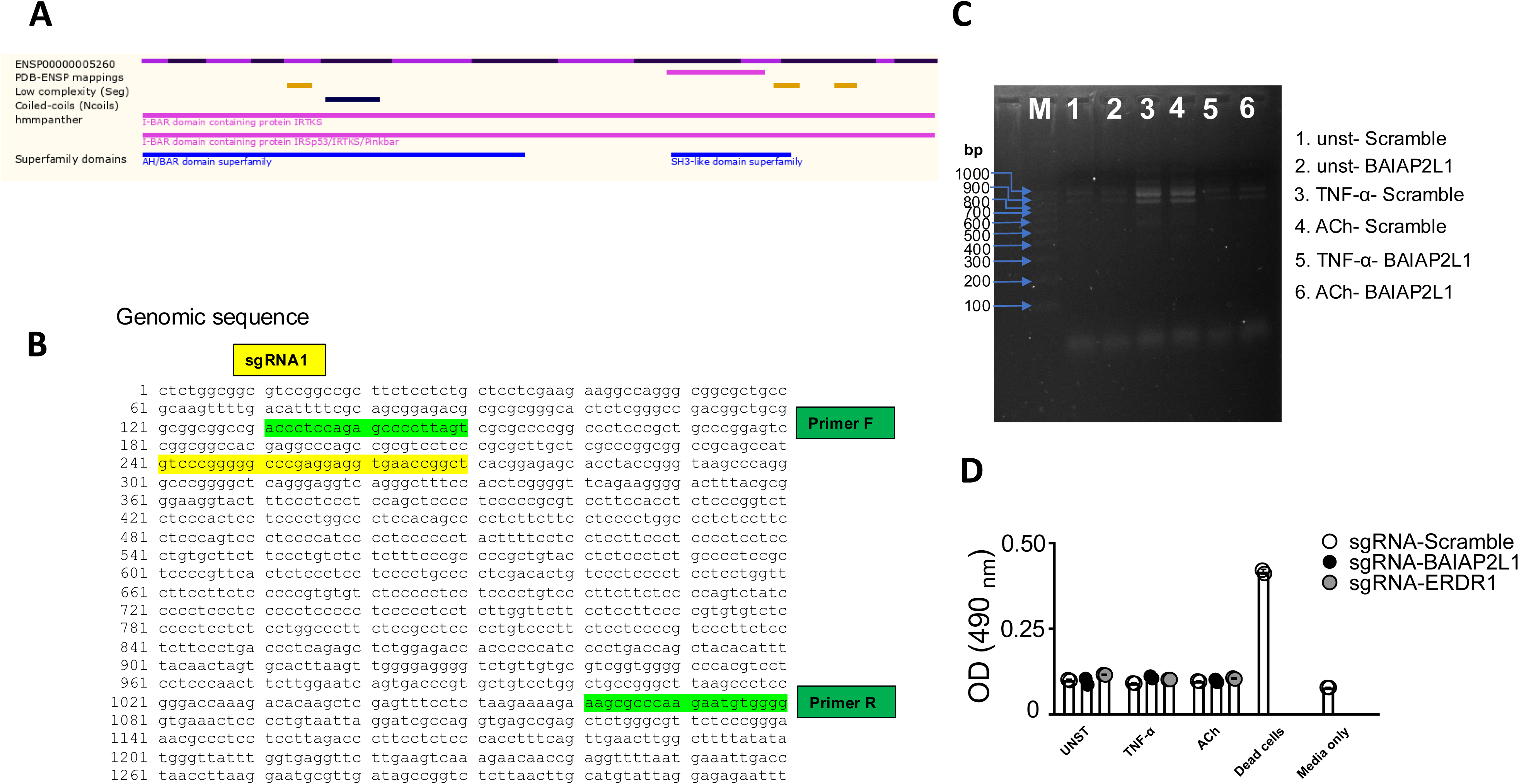
Targeting and validation of *BAIAP2L1* deletion by CRISPR. **(A)** Schematic of human *BAIAP2L1* coding region showing conserved N-terminal I-BAR domain and C-terminal SH3-like domain. **(B)** Genomic sequence of show target sgRNA1 region (yellow) and primer 2 regions (grey). **(C)** DNA gel (1.6%) from PCR showing DNA ladder and a 1000bp product in stimulated scramble transfected sgRNA (lanes 3-4) and reduced expression of the band in stimulated *BAIAP2L1* transfected sgRNA (lanes 5-6) and unstimulated cells (lanes 1-2) using primer 2 region for PCR. **(D)** Lactate dehydrogenase (LDH) assay showing absorbance at 490 nm in supernatants from sgRNA transfected cells treated with media alone, TNF-α or acetylcholine. Positive control are dead cells killed by 1% Triton X and control is media alone.

